# PROTRIDER: Protein abundance outlier detection from mass spectrometry-based proteomics data with a conditional autoencoder

**DOI:** 10.1101/2025.02.01.636024

**Authors:** Daniela Klaproth-Andrade, Ines F. Scheller, Georgios Tsitsiridis, Stefan Loipfinger, Christian Mertes, Dmitrii Smirnov, Holger Prokisch, Vicente A. Yépez, Julien Gagneur

## Abstract

**Motivation:** Detection of gene regulatory aberrations enhances our ability to interpret the impact of inherited and acquired genetic variation for rare disease diagnostics and tumor characterization. While numerous methods for calling RNA expression outliers from RNA-sequencing data have been proposed, the establishment of protein expression outliers from mass spectrometry data is lacking.

**Results:** Here, we propose and assess various modeling approaches to call protein expression outliers across three datasets from rare disease diagnostics and oncology. We use as independent evidence the enrichment for outlier calls in matched RNA-seq samples and the enrichment for rare variants likely disrupting protein expression. We show that controlling for hidden confounders and technical covariates, while simultaneously modeling the occurrence of missing values, is largely beneficial and can be achieved using conditional autoencoders. Moreover, we find that the differences between experimental and fitted log-transformed intensities by such models exhibit heavy tails that are poorly captured with the Gaussian distribution and report stronger statistical calibration when instead using the Student’s t-distribution. Our resulting method, PROTRIDER, outperformed baseline approaches based on raw log-intensities Z-scores or on differential expression analysis with limma. The application of PROTRIDER reveals significant enrichments of AlphaMissense pathogenic variants in protein expression outliers. Overall, PROTRIDER provides a method to confidently identify aberrantly expressed proteins applicable to rare disease diagnostics and cancer proteomics.

**Availability and Implementation:** PROTRIDER is freely available at <u>github.com/gagneurlab/PROTRIDER</u> and also available on Zenodo under the DOI zenodo.<u>15569781</u>.

**Contact:** Julien Gagneur: gagneur at in.tum.de

## Introduction

The detection of outliers in omics data, i.e. values that significantly deviate from the population and can thus be suggestive of a disease-causal gene, is of great importance for rare disease diagnostics (Kremer et al. 2017, Cummings et al. 2017, Yépez et al. 2022, Smail and Montgomery 2024). Importantly, outlier detection in omics data complements genome sequencing data by providing a functional readout to variants of uncertain significance whose interpretation is otherwise inconclusive. Outlier detection methods have been established for RNA-seq abundance, splicing, and DNA accessibility (Brechtmann et al. 2018, Mertes et al. 2021, Scheller et al. 2023, Çelik et al. 2024, Salkovic et al. 2023, Segers et al. 2023, Jenkinson et al. 2020, Labory et al. 2022, Salkovic et al. 2020). However, DNA accessibility and RNA sequencing cannot capture the effects of all pathogenic variants. Some variants may affect translation or protein stability, without impacting DNA accessibility or gene expression. To capture those effects, mass spectrometry-based proteomics constitutes an avenue to probe protein abundances as additional functional evidence (Kopajtich et al. 2021, Vialle et al. 2022, Hock et al. 2024). The interest in calling protein expression outliers also extends to cancer research, to characterize alterations in different molecular levels, find biomarkers, and explain drug sensitivities (Frejno et al. 2020, Roumeliotis et al. 2017).

Several studies have shown that measurements of gene expression, splicing, and chromatin accessibility data exhibit covariation patterns driven by biological and technical factors such as tissue, sampling site within the body, sex, batch, sequencing center, cause of death, sequencer, age, and read length (Kremer et al. 2017, Frésard et al. 2019, Mertes et al. 2021, Yépez et al. 2021, Çelik et al. 2024). Across those modalities, adjusting for these sources of covariation is strongly beneficial to enrich for the direct regulatory effects of genetic variants. Biological and technical sources of covariation also pertain to labeled proteomics experiments. Notably, samples analyzed together in the same batch of the mass spectrometry run exhibit a stronger correlation than those from different batches, especially for tandem mass tag labeled quantitative proteomics (Brenes et al. 2019, Zecha et al. 2019, Phua, Lim and Goh 2022). In a previous study, we proposed calling protein level outliers using a conditional autoencoder to account for hidden confounders and reported improvements over methods lacking this adjustment (Kopajtich et al. 2021).

Here we expand on and strengthen our previous work and present PROTRIDER. Method-wise, we investigate an alternative strategy to obtain the optimal encoding dimension, model the occurrence of missing values, compare linear against non-linear autoencoders, and perform a statistical assessment based on the Student’s t-distribution against the Gaussian distribution. Furthermore, we expand the benchmark to two other proteomics datasets of tumor cell lines and to enrichment among expression outliers in matched RNA-seq samples. Finally, we investigate the genetic determinants of the detected aberrant protein abundances, revealing that genes exhibiting protein expression outliers are strongly enriched for missense variants predicted to be pathogenic by AlphaMissense (Cheng et al. 2023).

## Methods

### Datasets

#### Mitochondrial disorder dataset

We used a dataset of 143 tandem mass tag (TMT) labeled quantitative proteomics samples with matched RNA-seq samples and variant calls from whole exome sequencing of individuals affected with a rare mitochondrial disorder of suspected genetic origin (Kopajtich et al. 2021). This dataset consisted of samples from patient-derived fibroblast cell lines by using a TMT 10-plex labeling reagent. Each TMT batch included 8 patient samples and 2 reference samples. The 143 samples were split over 21 TMT batches, with each batch contributing between 5 and 8 samples, except for one that only contributed one sample. Protein intensities were obtained from protein groups after peptide identification using MaxQuant v. 1.6.3.4 (Tyanova, Temu and Cox 2016). The RNA-seq samples were derived from the same fibroblast cultures and reads were counted using DROP (Yépez et al. 2021) as previously described (Yépez et al. 2022).

#### Tumor cell line panels

We additionally used proteomics measurements of the two publicly available tumor cell line panels NCI60 (n=60) and CRC65 (n=65), obtained from (Frejno et al. 2020). Variant calls from whole exome sequencing from the NCI60 cell lines were downloaded from CellMiner (<u>discover.nci.nih.gov/cellminer/</u>) in the form of the ‘DNA: Exome Seq - none’ processed dataset. For the CRC65 panel, somatic mutation calls from WES were only available for a subset of 33 of the cell lines through the DepMap project (depmap.org/portal/). We downloaded the ‘OmicsSomaticMutationsProfile.csv’ file containing the somatic variant calls, and the files ‘OmicsProfiles.csv’ and ‘Model.csv’ to map from the DepMap profile IDs to the cell line names of the CRC65 data in the proteomics data (Frejno et al. 2020). The variant calls for NCI60 were based on the hg19 genome build and annotated with ANNOVAR (Abaan et al. 2013, Wang, Li and Hakonarson 2010), whereas the variants obtained through DepMap were based on the hg38 genome build and annotated with VEP (v. 100.1) among other tools (DepMap 24Q2 Public 2024).

#### Proteomics data preprocessing

TMT-reference samples were excluded in this analysis and the remaining samples were not normalised using any reference samples. Raw protein intensities were log-transformed and adjusted for overall sample intensity using the DESeq2 size factor normalization (Love, Huber and Anders 2014), resulting in a protein intensity matrix **X** with elements sample *x_j,*i*_* for sample *i* and protein *j*.

### Aberrant protein expression level analysis with PROTRIDER

#### Conditional autoencoder

PROTRIDER uses an autoencoder to capture known and unknown sources of protein intensity variations, yielding expected log-intensities for each protein in each sample. Deviations of the measurements from these expected values are analyzed to identify outliers. First, proteins with more than a defined threshold of missing values across samples were filtered out. That threshold was set to 30% by default and other thresholds were further investigated. The remaining missing values were set to the protein-wise means in the input matrix **X** of the autoencoder and were ignored during mean squared error loss computation. We also considered as possible further input of the autoencoder the binary missingness mask **M,** in which ones indicate non-missing intensity values and zeros indicate missing values. In this setting, the intensity matrix **X** and the missingness mask **M** were stacked and jointly fed into the autoencoder, thereby modeling both protein intensities and missing value occurrences simultaneously. We also introduced the option to use a conditional autoencoder approach that explicitly uses specified covariates by including them both in the input of the encoder and the decoder.

The dimension *q* of the autoencoder bottleneck layer, i.e. latent space, was treated as a hyperparameter and optimized separately. We considered different numbers of layers for the encoder and the decoder, ranging between 1 and 3, where ReLU was used as an activation function between layers of the encoder and decoder, respectively. No non-linear activation was included for 1-layer encoders and decoders, effectively having a linear autoencoder of dimension *q*. In this case, the model was initialized with truncated Singular Value Decomposition after centering the protein intensity matrix **X** protein-wise and with bias terms adjusting for protein-wise means.

The model weights were optimized by minimizing a composite loss function consisting of two terms: (1) the mean squared error (MSE) between the predicted and observed protein intensities **X**, computed over all observed values, and (2) the binary cross-entropy (BCE) loss between the predicted probabilities of being observed and the missingness mask **M**. These two terms were combined as a weighted sum, with a predefined weighting parameter controlling the relative contribution of each.

The numerical optimization was performed with Adam (Kingma and Ba 2017) for 400 epochs. For 1-layer autoencoders, a small learning rate of 10^-4^ was used, whereas higher learning rates between 10^-4^ and 10^-3^ were used for multi-layer autoencoders.

#### Tail probability computation

For each sample *i* and protein *j*, the extremeness of the observed pre-processed intensity *x_i,j_* relative to the predicted intensity 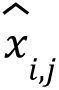 modeled by the autoencoder was quantified using two-sided tail probabilities, denoted *p_i,j_* To this end, we considered the residuals of the model *e_i,j_* defined as 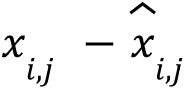 and computed their protein-wise means *m_j_* and unbiased standard deviations *s_j_*. The tail probabilities were obtained from either Gaussian or Student’s t-distribution. Gaussian tail probabilities were computed according to

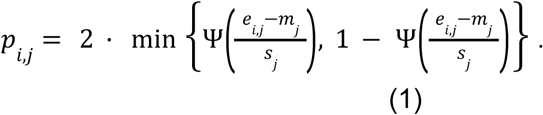

where ψ denotes the cumulative function of the normal distribution.

Early investigations with fitting Student’s t-distributions with protein-specific degrees of freedom yielded poor statistical calibration, probably due to numerical instability of the likelihood function with respect to the degree of freedom. Therefore, tail probabilities based on a Student’s t-distribution were robustly computed with a two-pass approach. In the first pass, we estimated the degrees of freedom, location, and scale parameters of the Student’s t-distribution using maximum likelihood for each protein. In the second pass, we set the degree of freedom for all proteins to a common value 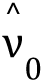 defined as the median of the degrees of freedom estimated in the first pass, and we fitted the location and the scale for each protein again. The two-sided tail probabilities, simply referred to as tail probabilities later on, were calculated as

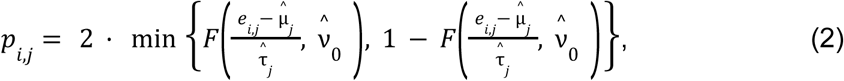

where *F* denotes the cumulative function of the Student’s t-distribution, 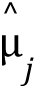 the location estimate, and 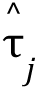 the scale estimates of protein *j*.

Tail probabilities were used to rank outlier candidates. No tail probabilities were reported for missing values.

#### Selection of the optimal encoding dimension

To find the optimal encoding dimension *q* of the autoencoder, i.e. the dimension of the autoencoder’s latent space, we employed two strategies: i) the optimal hard threshold (OHT) method (Gavish and Donoho 2014), which applies to the linear autoencoders without covariates only, and ii) a grid search over different values of *q*.

For the latter approach, at most 25 candidate values or up to half the sample size, whichever is smaller, are explored for finding *q.* These values are logarithmically spaced between 4 and half the sample size. For each candidate value, we fit the autoencoder after injecting the original dataset with artificial outliers generated with a frequency of 1 per 1,000 under a simulation scheme described earlier (Brechtmann et al. 2018). Specifically, the outlier intensity *x^o^_i,j_* for sample *i* and protein *j* was generated by shifting the observed preprocessed intensity *x_i,j_* by *z_i,j_* times the standard deviation *s_j_* of *x_i,j_*. The absolute value *z_i,j_* was drawn from a log-normal distribution with the mean of the logarithm equal to 3 and the standard deviation of the logarithm equal to 1.6, and with the sign of *z_i,j_* either positive or negative, drawn with equal probability:

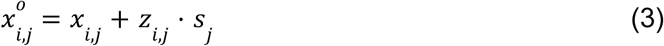

We select the candidate value for *q* that leads to the highest area under the precision-recall curve (AUPRC) of recovering the previously injected outliers when ranking by tail probabilities. After the optimal encoding dimension is determined either with the OHT or the grid search approach, the autoencoder is fitted using the determined value of *q* on the actual data without any artificially injected outliers.

#### Implementation

The autoencoder model of PROTRIDER was implemented in Python (v.3.8.13) using PyTorch (v.1.13.1). It is available at github.com/gagneurlab/PROTRIDER. The package includes the Python-based autoencoder implementation, calculates tail probabilities, and produces results tables. We also provide an example dataset and usage guidelines. The code is also available on Zenodo under the DOI zenodo.<u>15569781</u> (Klaproth-Andrade et al. 2025).

### False Discovery Rate assessment

#### False Discovery Rate definition

Akin to the multiple hypothesis testing problem for *P*-values, a nominal probability cutoff on tail probabilities would lead to a number of calls increasing with the number of proteins even in the absence of genuine outliers. We addressed this issue by defining the False Discovery Rate (FDR) in the context of our method. To this end, we considered a null model, in which the observed values are independently drawn from a Student’s t-distribution with parameters estimated from a PROTRIDER fit to a dataset. Any discovery (outlier call) from data generated from the null model is a false positive. Akin to Benjamini and Hochberg (Benjamini and Hochberg 1995), we defined the false positive proportion *Q* to be the proportion of false discoveries among the discoveries. We also defined *Q* to be equal to 0 if no discovery is made. We defined the FDR as the expected value of *Q*.

#### Empirical FDR control assessment

We assessed whether the Benjamini-Yekutieli (Benjamini and Yekutieli 2001) and the Benjamini-Hochberg (Benjamini and Hochberg 1995) procedures applied to the tail probabilities (instead of *P*-values) controlled the above-defined FDR.

To this end, we generated data under the null model. Specifically, we sampled residuals e*_*i,j*_ for all samples *i* and proteins *j*, from a Student’s t-distribution with location, scale, and degrees of freedom parameters estimated on the mitochondrial disorder dataset from the residuals *e_i,j_* as described in the Tail probability computation subsection. The sampled residuals *e*_i,j_* were added to the PROTRIDER fit 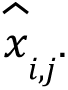 We then reversed the original preprocessing transformations to yield a synthetic protein intensity matrix with no true outliers.

The entire PROTRIDER fitting procedure was applied to each of 100 simulated null datasets. Next, the Benjamini and Hochberg procedure and the Benjamini and Yekutieli procedure were applied to the tail probability vectors of each sample from each simulated dataset, and outlier calls were made at a fixed threshold. Since the datasets were simulated under the null model, any detected outlier constituted a false positive, yielding false discovery proportions in each run equal to 0 (no calls made) or 1 (some calls made). The empirical FDR was estimated as the average false discovery proportion across the 100 simulations and across all samples. Moreover, the total amount of calls was recorded for each simulation and sample combination.

#### Final set of outliers with controlled FDR

To define a set of final protein-sample outliers on the mitochondrial disorder dataset, as well as on the tumor cell line panels, while controlling for the FDR, we applied the procedure of Benjamini and Yekutieli, and alternatively, the one of Benjamini and Hochberg sample-wise, providing tail probabilities instead of *P*-values. Protein outliers were defined as those with an FDR of 0.1 or lower.

### Cross-validation

We also ran PROTRIDER in a leave-one-out and in a 5-fold cross-validation setting. In the leave-one-out setting, each sample was sequentially held out as a test case while the remaining samples were used for model training and hyperparameter tuning. The optimal encoding dimension was determined on the training set using both grid search and OHT approaches. During autoencoder training, 20% of the training set was set aside as a validation set for early stopping. After training, residuals were obtained for all training samples, and for each protein, a Student’s t-distribution was fitted in a two-pass approach as described above. The resulting model was then applied to the held-out test sample to derive tail probabilities. This process was systematically repeated for every sample in the dataset.

In the 5-fold setting, the dataset was split into 5 folds. Each fold was used once as a test set while the remaining folds were used for training. The tail probabilities for each test fold were derived using the same procedure as in the leave-one-out setting.

### Limma

We used the differential expression detection method limma (Ritchie et al. 2015) to call outliers by testing each sample individually against all other samples. To run limma, we used the preprocessed protein intensities *x_i,j_* as the input. Relevant covariates for each dataset were included through limma’s design matrix. This resulted in a different limma run for each sample of each dataset. The results were concatenated per dataset into a table consisting of multiple testing-corrected *P*-values, fold-changes, and z-scores for each sample-protein combination.

### Z-score

We computed Z-scores from the preprocessed protein intensities *x_i,j_* by subtracting the mean and dividing by the unbiased standard deviation protein-wise. Tail probabilities were calculated using the normal distribution, and the procedure of Benjamini and Yekutieli (Benjamini and Yekutieli 2001) was applied sample-wise.

### Isolation forest

As an alternative method, we also called outliers by fitting a protein-specific Isolation forest (Liu, Ting and Zhou 2012) model in two configurations: (1) fitted directly on the preprocessed intensities *x_i,j_*, and (2) fitted on the residuals *e_i,j_* returned by PROTRIDER. Outlier candidates were ranked by the anomaly scores predicted by the Isolation forest models.

### Variant annotations

To evaluate PROTRIDER on independent benchmarks, we considered different types of rare variant categories. Variants were considered rare if they had a minor allele frequency of 0.1% or lower according to gnomAD v2.1.1 (Karczewski et al. 2020). Variants on the mitochondrial disorder dataset were annotated using the consequences from the Variant Effect Predictor VEP v99 (McLaren et al. 2016), AbSplice (Wagner et al. 2023), AlphaMissense (Cheng et al. 2023), and CADD v1.6 (Rentzsch et al. 2019). For VEP annotations, when more than one rare variant was present in a gene for a given sample, we considered only the most severe consequence based on Ensembl’s severity ranking (https://www.ensembl.org/info/genome/variation/prediction/predicted_data.html).

AlphaMissense classifications were only considered for gene-sample combinations without any variant categorized as high-impact by VEP in the same gene and replaced the VEP missense annotations.

On the tumor cell line datasets, we used the variant annotations containing ANNOVAR consequences for the NCI60 samples and VEP annotations for the CRC65 samples. Additionally, we annotated AlphaMissense predictions in both datasets. As the variant calls corresponding to the tumor cell lines were somatic variant calls, we considered only variants with a sample allele frequency of at least 50%.

### Enrichment for rare variants likely disrupting protein abundance

To benchmark underexpression outlier calls, we considered three non-disjoint sets of rare variants that could disrupt protein abundance: i) stop, frameshift, direct splice-site, and missense variants annotated by VEP, ii) likely deleterious variants based on CADD with a CADD PHRED-scaled score of at least 20, and iii) variants predicted as pathogenic by AlphaMissense. For each set, proportions were computed among all underexpression outliers.

### Aberrant expression analysis of RNA-seq samples

To compare the protein abundance outlier calls to RNA expression outliers in the mitochondrial disorder dataset, we ran the gene expression outlier detection method OUTRIDER (Brechtmann et al. 2018) as part of the Detection of RNA Outlier Pipeline DROP v1.0.3 (Yépez et al. 2021). We used the GRCh37 primary assembly, release 29, of the GENCODE project as the reference genome (Frankish et al. 2019). RNA expression outliers were defined as those with an FDR of 0.1 or lower.

## Results

We developed PROTRIDER, a model to call aberrant protein expression from mass spectrometry-based quantitative proteomics data. PROTRIDER models log-transformed protein intensities adjusted for overall sample intensity using a conditional autoencoder to account for biological and technical sources of variation, which may be known or unknown (Fig. 1). Known covariates such as batch effects, which are often strongly observed in TMT-labeled proteomics data (Brenes et al. 2019), can be directly provided to the model.

**Fig. 1:**
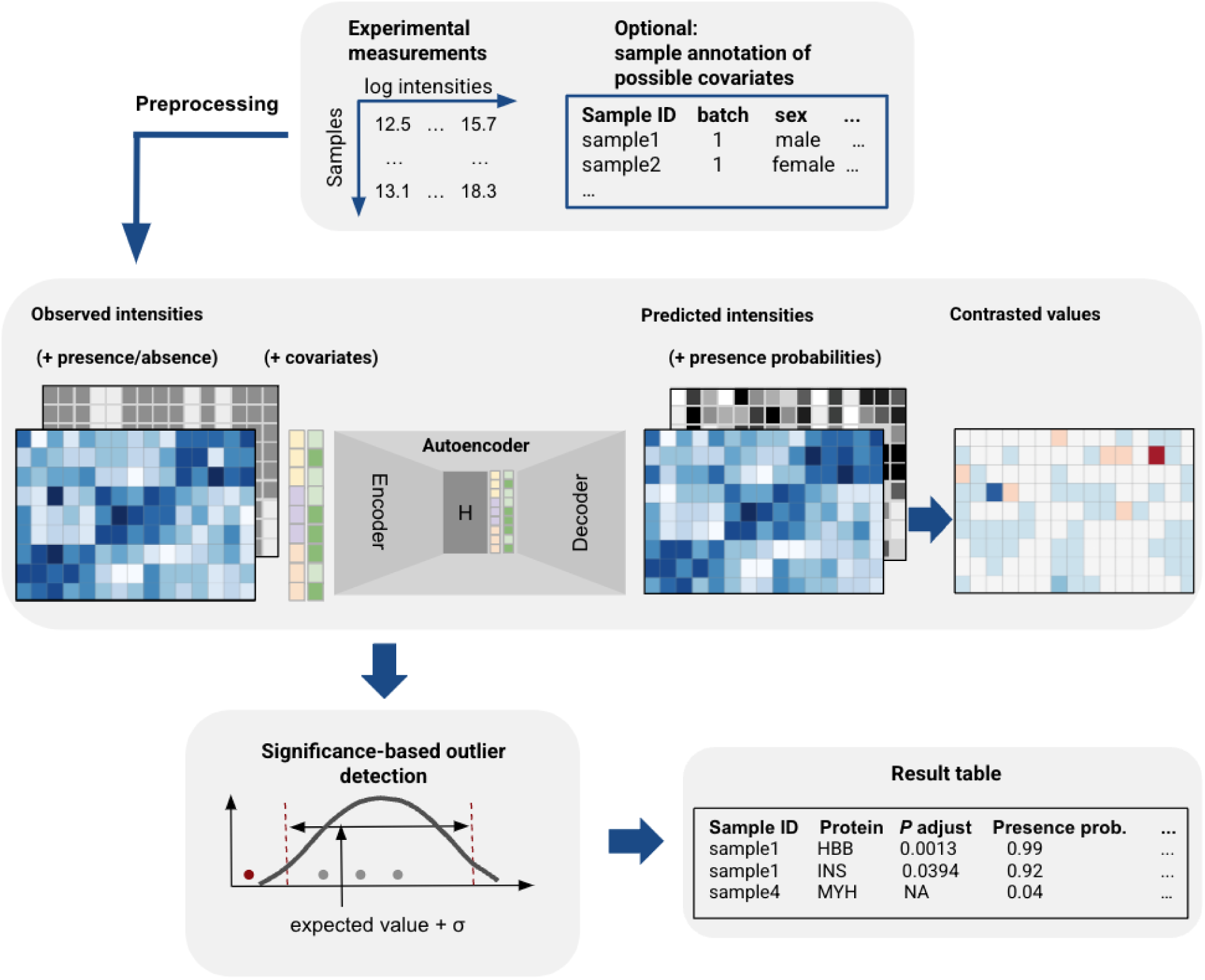
Schematic overview of the PROTRIDER outlier detection approach. PROTRIDER takes a protein intensity matrix from a quantitative proteomics experiment as input, and, optionally a missingness mask, as well as covariates such as batch and sex, and fits a conditional autoencoder to account for known and unknown biological and technical sources of covariation of proteins across samples. Expected log-transformed protein intensities from the autoencoder are then contrasted with the observed values and tested for statistical significance.

Proteomics data often contain missing values, which complicates model fitting (Brenes et al. 2019, Phua, Lim and Goh 2022). To alleviate the impact of missing values, we filtered out proteins with more than 30% missing values across an entire dataset. The remaining missing values were ignored in the mean squared error loss computation and tail probability calculations. Moreover, we considered modeling the occurrences of missing values jointly with the observed intensities in the autoencoder (Methods).

To identify the optimal latent space dimension, we employed two different approaches. The first one conducts a grid search across candidate values for the encoding dimension and selects the one that optimizes the recovery of artificially injected outliers (Methods). The second option applies to linear autoencoders without covariates and without missingness modelling only and is based on the Optimal Hard Threshold (OHT) procedure, an analytical solution to the number of principal components to select when assuming the data matrix sums to a low-rank matrix and a white noise matrix (Gavish and Donoho 2014). OHT has been recently used as a computationally efficient alternative to the grid search approach in the context of RNA-seq outlier calling by the OutSingle and saseR methods (Salkovic et al. 2023, Segers et al. 2023).

After the autoencoder model of PROTRIDER is fitted, the residuals, i.e., the differences between the observations and the autoencoder predictions, are used to compute two-sided tail probabilities assuming either a normal distribution or a Student’s t-distribution fitted for each protein.

We applied PROTRIDER to three datasets. The first dataset comprises 143 TMT-labelled proteomics samples collected from individuals with a rare, suspected Mendelian, mitochondrial disorder, which we refer to as the mitochondrial disorder dataset (Kopajtich et al. 2021). After filtering out proteins with more than 30% missing values across samples, 7,060 proteins were quantified (Supplementary Fig. S1). Additionally, we considered proteomic measurements of two tumor cell line panels, NCI60 and CRC65 (Frejno et al. 2020). The NCI60 panel contains 60 cell lines from various tissues, while the CRC65 panel consists of 65 colorectal tumor cell lines. The two panels were analyzed separately because they differed in the number of proteins detected, tumor entities present in the data, genome build, and exome sequencing processing tool. After removing proteins with more than 30% missing values, the NCI60 dataset was left with 6,755 proteins and the CRC65 dataset with 8,430 proteins (Supplementary Fig. S2).

Strong batch effects were observed in the raw log-transformed protein intensities between samples for all three datasets (Fig. 2A, Supplementary Fig. S3A, C). In the mitochondrial disorder dataset, they were largely driven by the different TMT batches (Fig. 2A). Similarly, in both of the cell line panels, strong correlations between samples were observed in the experimental measurements, driven by the tissue of origin in NCI60 and MSI (Microsatellite Instability Biomarker) status, and subtype classification in CRC65 (Supplementary Fig. S3A, C).

**Fig. 2:**
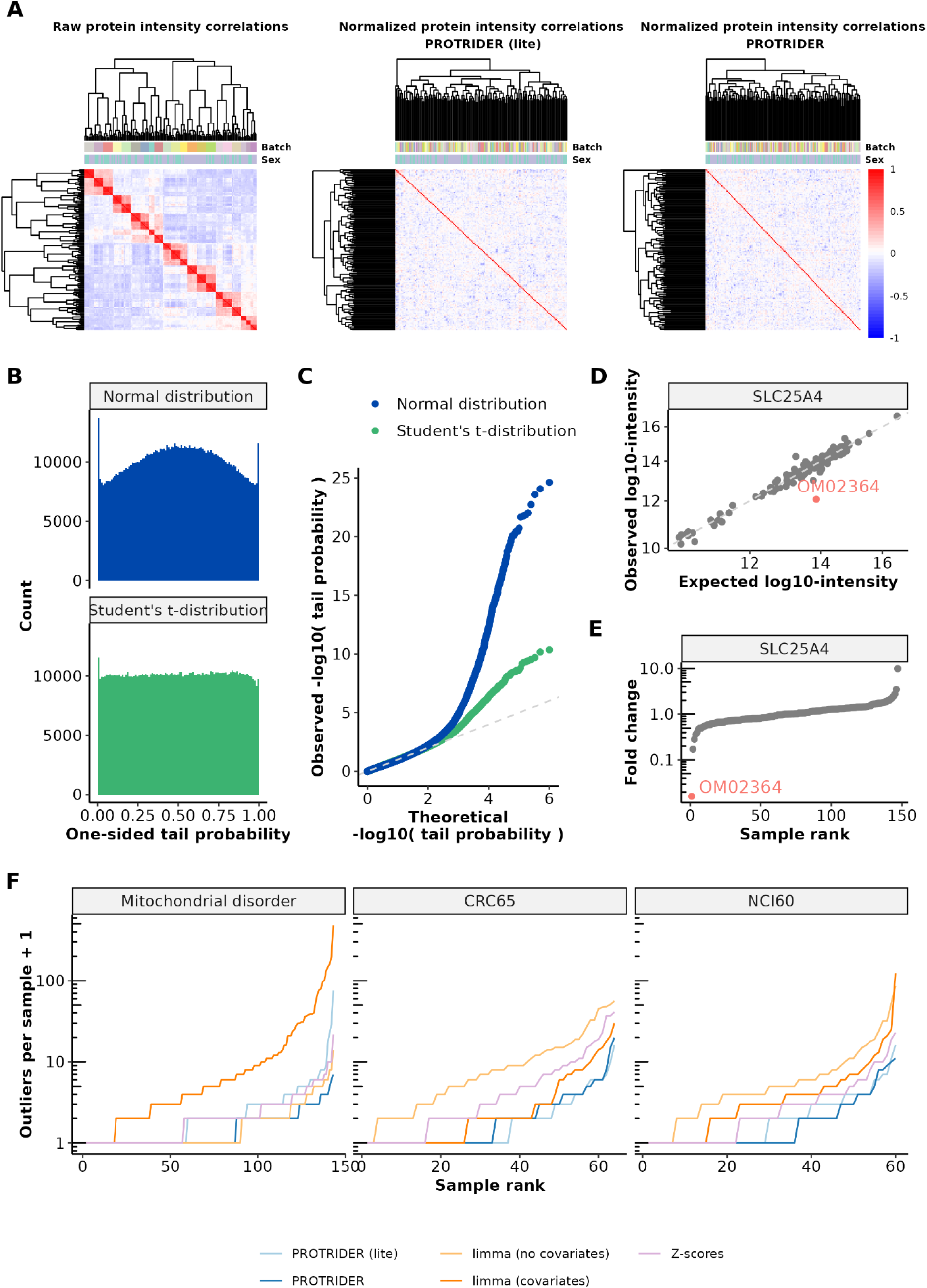
PROTRIDER accounts for known and hidden covariates and typically detects a limited number of outliers per sample. **A)** Heatmaps of sample-sample correlations of protein log-transformed intensities on the mitochondrial disorder dataset before (left) and after the PROTRIDER with OHT (middle) and with autoencoder correction using sex and batch as covariates (right). **B)** Histogram of tail probabilities from the normal distribution (blue) and from the Student’s t-distribution (green) obtained after the PROTRIDER (lite) correction on the mitochondrial disorder dataset. **C)** Quantile-Quantile plot comparing observed-log_10_(tail probabilities) to their theoretical quantiles under the null (uniform distribution). **D)** Observed against expected log-transformed protein intensities for the gene SLC25A4, highlighting the protein underexpression outlier on individual OM02364. **E)** Sorted fold changes of protein intensities (ratio of observed and fitted values) for the gene SLC25A4, highlighting individual OM02364. **F)** Sorted number of protein abundance outliers per sample obtained by the four methods: PROTRIDER (dark blue), PROTRIDER (lite, light blue), the limma-based approaches with (dark orange) and without covariates (light orange), and the Z-score-based (violet) approach on the three datasets (facets).

To account for such sources of covariations, we evaluated the effect of explicitly fitting a conditional autoencoder model that received the covariates directly as part of its input, in addition to the protein intensities, as well as missingness modelling. We refer to this model as ‘PROTRIDER’. We compared it to an autoencoder that did not have access to information about potential covariates or the missingness mask during model fitting. To this end, we use a linear autoencoder that we initialized using truncated singular value decomposition and with a dimension determined with the OHT approach. The parameters of this model were further tuned (Methods). We named this model ‘PROTRIDER (lite)’. On the mitochondrial disorder dataset, we included sex, TMT-batch, sample preparation batch, and sequencing instrument of each sample as covariates, whereas we included sex, age, and tissue of origin for NCI60, and MSI status and subtype classification for CRC65. We found encoding dimensions of 59, 13, and 18 to be optimal for PROTRIDER on the mitochondrial disorder dataset, NCI60, and CRC65, respectively. For PROTRIDER (lite), encoding dimensions of 40, 7, and 5 were obtained, respectively. Notably, we observed that the relationship between the encoding dimension and the performance in recovering artificially injected outliers was typically well-behaved and unimodal, with a relatively wide range of near-optimal values for the encoding dimension (plateau, Methods, Supplementary Fig. S4).

Both PROTRIDER versions were able to account for the observed strong batch effects. Specifically, the normalized intensities after fitting the autoencoder models were no longer strongly correlated between samples (Fig. 2A). On the mitochondrial disorder dataset, the median within-batch pairwise sample Spearman correlations were reduced from 0.83 ± 0.059 (mean ± standard deviation, here and elsewhere) to 0.15 ± 0.087 for PROTRIDER (lite) and to 0.13 ± 0.079 for PROTRIDER. These results show that despite not having access to covariates, PROTRIDER (lite) succeeded in capturing the covariation. Similar results were obtained on the tumor cell line panel datasets (Supplementary Fig. S3B, D).

When modeling the residuals with protein-specific normal distributions, we observed an excess of one-sided tail probabilities close to 0.5, as well as close to 0 and 1, indicative of the data exhibiting heavy tails (Fig. 2B). Therefore, we opted to use the Student’s t-distribution instead of the normal distribution. Fitting a Student’s t-distribution for each protein was robustly achieved by learning a value for the degrees of freedom common to all proteins (Methods). The tail probability distributions (observed via histograms and quantile-quantile plots) indicated that the Student’s t-distribution yielded much better statistical calibration than the normal distribution (Fig. 2B, C).

PROTRIDER does not perform hypothesis testing and therefore does not return p-values. Instead, it reports tail probabilities of the model fitted to the data. Thus, to formalize the issue of false discoveries in this context, we considered a null model in which the observed values are independently drawn from a Student’s t-distribution with parameters estimated from a PROTRIDER fit to a dataset. Any discovery (outlier call) from data generated from the null model is a false positive. We confirmed that the tail probabilities under this null model were approximately uniformly distributed when fitting from scratch PROTRIDER models to 100 such simulated datasets (Methods, Supplementary Fig. S5A,B).

Next, we defined the false discovery rate (FDR) as the expected proportion of false discoveries among discoveries, akin to Benjamini and Hochberg (Benjamini and Hochberg 1995) in the context of hypothesis testing (Methods). We then applied the Benjamini-Yekutieli (BY, (Benjamini and Yekutieli 2001)) procedure and the Benjamin-Hochberg (BH, (Benjamini and Hochberg 1995)) procedure to the tail probabilities, each in a sample-wise manner. Our empirical evaluation showed that both procedures successfully controlled the FDR across different thresholds (Supplementary Fig. S5C) for data simulated under the null model, with a limited number of false positives observed at a threshold of 0.1 (Supplementary Fig. S5D). The BY procedure is more general than the BH procedure as it can be applied to dependent statistics. Consequently, it yielded a more conservative control of the FDR on simulations compared to BH (Supplementary Fig. S5C). However, we note that the empirical calibration of the BH procedure might be optimistic because the simulations were performed using independently drawn errors. Therefore, we conservatively used the BY procedure for the subsequent analyses (FDR < 0.1).

We exemplify PROTRIDER outlier calls for a rare disease diagnostic relevant case we reported earlier (Kopajtich et al. 2021). The mitochondrial solute carrier SLC25A4, which translocates ADP from the cytoplasm into the mitochondrial matrix and ATP from the mitochondrial matrix into the cytoplasm, appears as a downregulated outlier in the individual OM02364 (FDR = 0.05). This is evident from the deviation of the observed intensity compared to the PROTRIDER expectation by 61-fold (Fig. 2D, E), which is aberrant compared to the variations seen across the other individuals. This aberrantly reduced expression of the mitochondrial disorder-causing protein SLC25A4 validated the functional impact of the heterozygous missense candidate variant from the patient and led to the patient’s genetic diagnostics (Kopajtich et al. 2021).

To provide baseline comparisons, we considered Z-scores computed from the observed log-transformed protein intensities adjusted for overall sample intensity (denoted as ‘Z-scores’). Moreover, we considered an approach based on the differential expression test limma (Ritchie et al. 2015), applied by testing each sample against all others. While limma is not specifically designed for outlier detection, it supports covariate adjustment. We reasoned that it can still serve as a useful baseline for comparison and offers context for understanding the performance of our method relative to a commonly accessible alternative.

At an FDR of 0.1, PROTRIDER and PROTRIDER (lite) typically reported 1 outlier per sample across all three datasets (Fig. 2F). This is in line with the number of gene expression outliers typically reported by OUTRIDER on RNA-seq samples (Yépez et al. 2022). In contrast, the limma approaches typically yielded larger and more variable numbers of outliers across the datasets (median up to 7 outliers per sample, depending on the dataset and the inclusion of covariates, Fig. 2F).

To benchmark alternative methods for protein outlier detection performance using orthogonal data, we first considered enrichments for variants that likely lead to protein abundance outliers. To this end, we selected variants that are rare in the human population with a minor allele frequency (MAF) less than or equal to 0.1% in gnomAD (Karczewski et al. 2020) and whose consequence is stop, frameshift, splice-site, or missense according to VEP, likely deleterious according to CADD, or pathogenic according to AlphaMissense (McLaren et al. 2016, Cheng et al. 2023, Rentzsch et al. 2019). Those variants do not necessarily lead to aberrant protein abundances. However, one can expect that more accurate protein abundance outlier callers will lead to higher enrichments for genes carrying those variants. We first compared linear autoencoders from PROTRIDER to the limma-based and Z-scores approaches, using the variant categories described above as a ground truth proxy. PROTRIDER and PROTRIDER (lite) outlier calls performed consistently favorably across all variant categories (Fig. 3A).

**Fig. 3:**
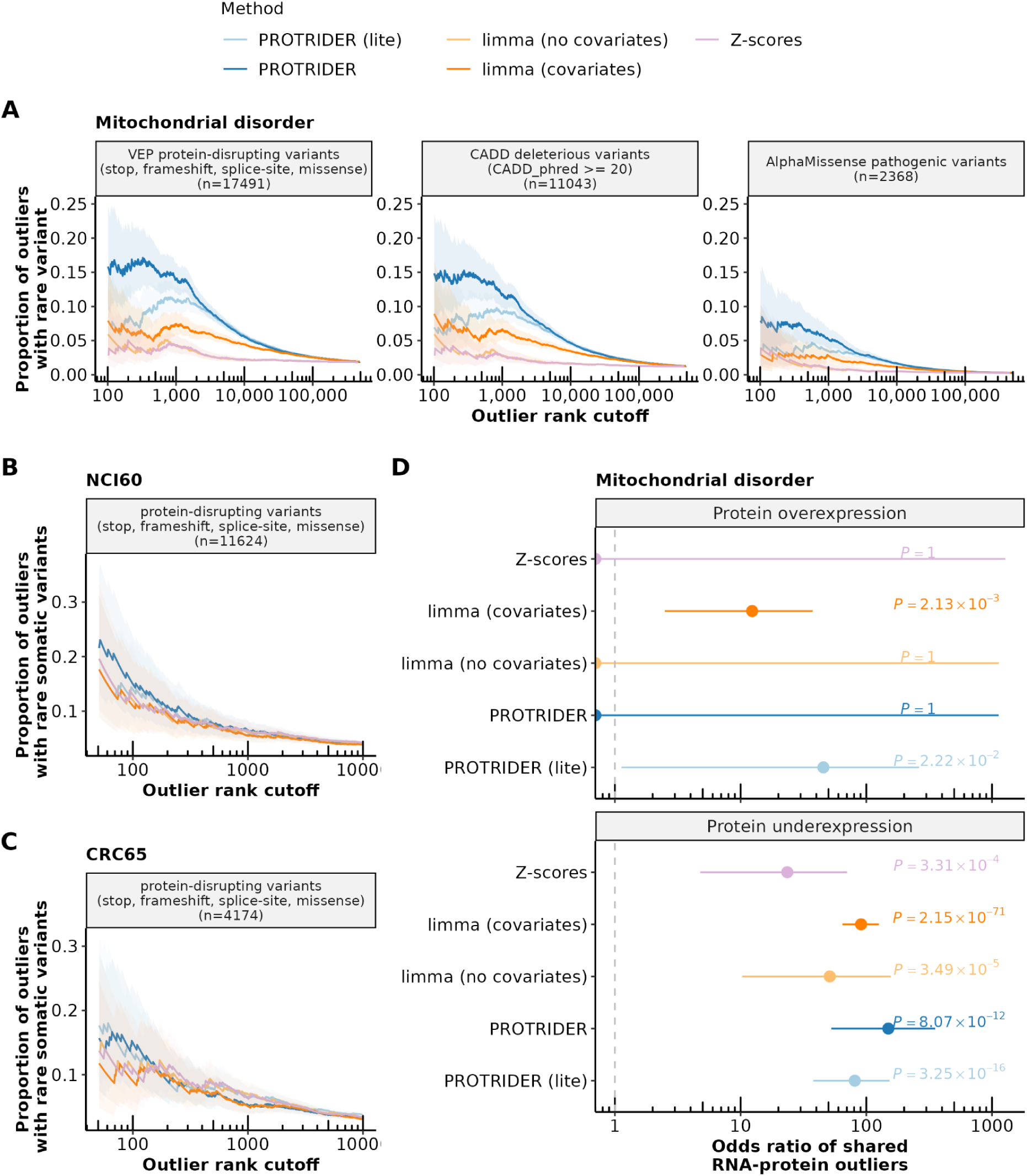
PROTRIDER outperforms baseline approaches on rare variant benchmarks. **A)** Proportion of outliers with at least one rare variant on the mitochondrial disorder dataset for underexpression outliers calls from PROTRIDER (blue), PROTRIDER (lite, light blue), the limma-based method with covariates (orange) and without covariates (light orange), and the Z-score-based method (violet) on three sets of rare variant categories as ground truth proxies: i) VEP stop, frameshift, direct splice-site, and missense variants, ii) CADD deleterious variants (PHRED score ≥ 20) and iii) AlphaMissense pathogenic variants. Ribbons mark 95% confidence intervals. **B-C)** Same as A) but for the two tumor cell line panels NCI60 and CRC65, and only on the category of stop, frameshift, direct split-site, and missense variants. **D)** Odds ratio of shared RNA and protein outliers in the mitochondrial disorder dataset and their 95% confidence intervals (Fisher’s test) for the two PROTRIDER methods, the two limma-based approaches, and the Z-scores approach.

Specifically, among the 1,000 top-ranked underexpression outliers detected by PROTRIDER, 14% ([11.9, 16.3] 95% confidence interval) of the called outliers harbored at least one rare variant whose consequence was stop, frameshift, splice-site, or missense according to VEP on the mitochondrial disorder dataset. In comparison, only 7.4% ([5.9, 9.2] 95% confidence interval) of the outliers called by the limma-based differential expression approach, including covariates harbored at least one such rare variant. The Z-score approach, along with a limma-based differential expression without covariates, showed substantially lower proportions (never exceeding 5%). The superiority of PROTRIDER was observed independently of the rank cutoff and for all three considered variant categories (Fig. 3A). Moreover, we compared the results obtained when filtering the dataset to include proteins with at most 30% missing values to those obtained using alternative thresholds for the maximum allowed proportion of missing values per protein. Across all thresholds, PROTRIDER consistently outperformed the other methods. There is a tradeoff between protein coverage and model performance, whereby a lower proportion of missing values yielded a higher proportion of underexpression outliers associated with rare variants likely disrupting protein expression, yet at the cost of excluding more proteins from the analysis, potentially omitting valuable outlier candidates (Supplementary Fig. S6). As a compromise, we opted for a threshold of at most 30% missing values per protein for all subsequent analyses.

For the two tumor cell line datasets, no significant difference between the methods was found when comparing them at equal protein ranks, and the proportions of rare variants likely disrupting protein expression were globally lower compared to the mitochondrial dataset. On possible explanation for the weaker proportions is the lower sample size (60 and 65 versus 143). Consistent with this hypothesis, the proportions of variants decreased gradually as we downsampled the mitochondrial dataset, yielding proportions at matched sample sizes that were similar to those observed in the tumor cell line dataset (n=60, Supplementary Fig. S7). Additionally, the lower performance observed in the tumor cell lines compared to the mitochondrial disorder dataset may also reflect greater genetic heterogeneity in tumor-derived cell lines relative to patient fibroblasts. We evaluated whether changing the process of artificially injecting outliers during the selection of the encoding dimension could improve the model. Specifically, changing the outlier injection mean from 3 to 6 consistently resulted in substantially higher AUPRC when recovering artificially injected outliers for all encoding dimension candidates (Supplementary Fig. S8a). However, even though the optimal dimensions changed for the two panels (from 13 to 8 for NCI60 and from 18 to 10 for CRC65), this did not practically result in a difference in the variant enrichment performance (Supplementary Fig. S8b,c).

Nonetheless, considering the significant calls only, PROTRIDER reported much fewer outliers than the limma-based approach on the tumor cell line panels (75 vs. 332 on NCI60, 104 vs. 194 on CRC65) and maintained in the case of NCI60 a reasonably high proportion of rare variants likely causing protein abundance aberration compared to the limma-based approach (8.8% vs 7.6% at rank 500, Fig. 3B, C). Altogether, this benchmark using independent evidence from rare genetic variants indicates that PROTRIDER in either setting improves the detection of genuinely aberrantly expressed proteins.

We also considered whether non-linear autoencoders could improve over linear ones. To this end, we added up to three layers with non-linear activations between layers to PROTRIDER. However, using multi-layer models did not improve the performance over the one-layer model on the mitochondrial disorder dataset (Supplementary Fig. S9). Therefore, we used one-layer autoencoders for all the subsequent analyses.

Moreover, we also considered a non-parametric approach to call outliers. To this end, we applied isolation-based anomaly detection using Isolation Forests (Liu, Ting and Zhou 2012). Since our goal is to identify outlier intensities per protein and sample, Isolation Forest was run on each protein individually. We first applied Isolation Forest directly on the preprocessed protein intensities. This led to essentially no enrichment for variants likely disrupting protein expression (Fig. S10). We next applied Isolation Forest on the residuals returned by PROTRIDER. This second approach outperformed limma with known covariates, confirming the importance of adjusting for hidden sources of covariation, yet did not improve upon PROTRIDER (Fig. S10). These results are consistent with the good calibration of PROTRIDER tail probabilities evident from the QQ-plot analysis and further indicate that t-distributions reasonably capture protein intensity residuals.

Furthermore, we investigated the contribution of various modeling choices by fitting PROTRIDER under different configurations (Supplementary Fig. S11). Our analysis showed that modeling missingness as well as using PCA for weight initialization improves PROTRIDER’s performance. Interestingly, PROTRIDER (lite) achieves similar performance regardless of whether PCA-based weight initialization is used. Comparable results to those obtained with PROTRIDER (lite) are also observed when using a PCA projection alone, without subsequent autoencoder training. We observed consistently similar performance across a range of weighting factors used to combine the binary cross-entropy and mean squared error loss terms when incorporating the binary missingness modelling, indicating that the model is relatively robust to this parameter choice (Supplementary Fig. S12).

We note that our approach does not require a cross-validation scheme because the encoding dimension is smaller than the dataset dimension, and because the encoding dimension is chosen to optimize the recovery of original, uncorrupted intensities. Nonetheless, we investigated whether a cross-validation scheme could further improve the robustness to PROTRIDER. Specifically, we considered both five-fold cross-validation and leave-one-out cross-validation (Methods). We observed lower performance for both approaches on the variant enrichment benchmark, whereby the leave-one-out approach performed substantially better than five-fold cross-validation (Supplementary Fig. S13). This consistent trend indicates that the model needs the entire dataset to provide good fits, at least for the sample sizes we could investigate.

As another independent type of benchmarking data, we considered gene expression outliers from RNA-sequencing. As for the rare variant benchmark performed above, this benchmark only provides a ground truth proxy, as some gene expression outliers can be translationally buffered. Moreover, some protein abundance outliers may not be reflected in RNA-seq, for instance, due to post-transcriptional regulation (Kusnadi et al. 2022, Wang et al. 2018). Nevertheless, the proportions of RNA-seq outliers obtained on the same samples can be used to compare different protein abundance outlier callers. Another added value of using RNA-seq outliers for benchmarking, compared to variants likely disrupting protein expression, is that they allow assessment of the overexpression outliers. All methods obtained a significant enrichment (Fisher’s test nominal *P*-value < 0.05) for underexpression RNA-seq outliers called by OUTRIDER (Brechtmann et al. 2018). However, the enrichment for the Z-scores approach was two times lower than for the other three methods, which performed similarly (Fig. 3D). For overexpression, no enrichment for RNA-seq outliers was found by the limma approach without covariates, nor by the Z-scores approach, nor by PROTRIDER, including missingness modelling, while the remaining methods performed similarly and significantly higher than random (Fig. 3D).

Taken together, these results show that PROTRIDER is well suited for the task of aberrant protein abundance detection, outperforming the Z-scores approach and limma with and without covariates on the benchmark using rare protein-disrupting variants and outperforming the limma approach without covariates in the enrichment of shared protein and RNA expression outliers.

Having established a protein expression outlier caller, we next investigated various characteristics of the genetic determinants of these outliers. To this end, we considered all rare variants (gnomAD MAF <0.1%) falling within the gene boundary. We note that, as the variants of this dataset were called from whole exome sequencing, they were biased toward the coding sequence. We found that 30% of the underexpression protein abundance outliers called by PROTRIDER had at least one such rare variant (Fig. 4A). As expected, stop and frameshift variants showed the strongest enrichments (Fig. 4B) and explained overall 15.3% of the underexpression outliers (Fig. 4A). PROTRIDER achieved stronger enrichments, whereas PROTRIDER (lite) showed similar overall patterns, albeit with somewhat reduced signal strength (Fig. 4A, B). No significant enrichments for these variants were found, as expected, for overexpression outliers (Supplementary Fig. S14).

**Fig. 4:**
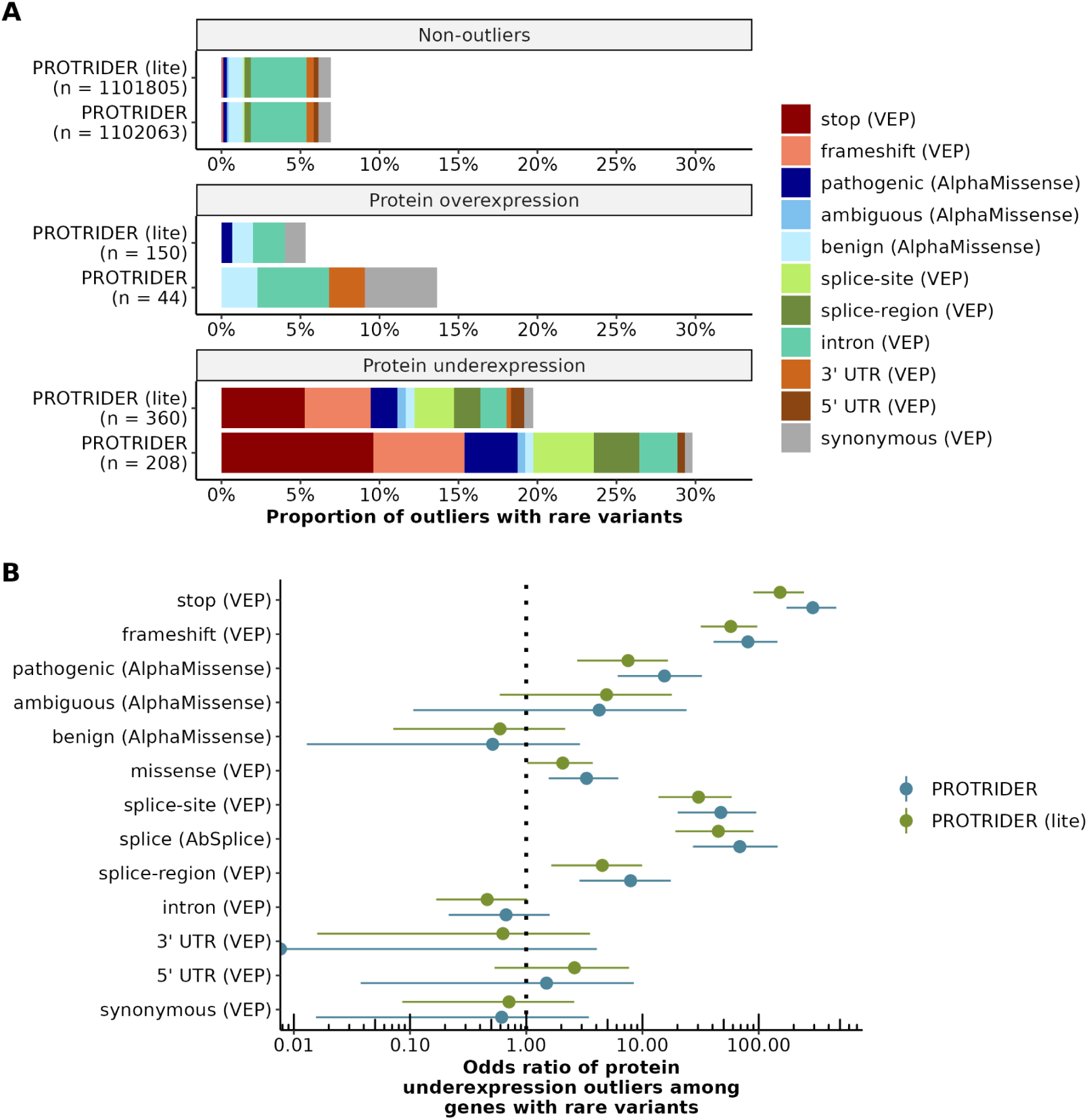
Genetic determinants of protein expression outliers. **A)** Proportions of non-outliers, protein overexpression, and protein underexpression outliers with a rare variant detected in the same gene for PROTRIDER and PROTRIDER (lite) in the mitochondrial disorder dataset. Colors indicate the different variant categories based on annotations from VEP and AlphaMissense. **B)** Odds ratios and their 95% confidence intervals (Fisher’s test) of the enrichment of the proportion for each variant category among underexpression outliers compared to the background proportion of the non-outliers for PROTRIDER (blue) and PROTRIDER (lite, green).

We further found a strong enrichment for AlphaMissense pathogenic variants (Fig. 4A, B), which explained another 3.4% of the underexpression outliers. The AlphaMissense categories helped prioritize missense variants affecting protein expression, as the AlphaMissense pathogenic category (odds ratio = 15.4) had stronger enrichment than the ambiguous (odds ratio = 4.2) and benign (odds ratio = 0.5) categories (Fig. 4B).

We evaluated the effect of replacing the conservative Benjamini-Yekutieli method with the Benjamini-Hochberg procedure for controlling the FDR. This substitution increased the number of outliers detected by about 69% (208 vs. 349) for PROTRIDER and about 134% (360 vs. 843) for PROTRIDER (lite), accompanied by only a slight decrease in the enrichment of protein-disrupting rare variants (Supplementary Fig. S15).

## Discussion

We described PROTRIDER, a model that extends the autoencoder-based outlier detection approach already valuable in the RNA-seq-based diagnosis of rare disease patients to mass spectrometry-based proteomics measurements. Using rare genomic variants and RNA-seq outliers as orthogonal data for benchmarking, we showed that protein abundance underexpression outliers detected with PROTRIDER outperformed baseline methods based on differential expression and simple Z-scores. We found that modeling the occurrence of missing values improved the model performance. In the three investigated datasets from rare disease and tumor cell lines, linear models outperformed non-linear, multi-layer autoencoders. Moreover, we found that the heavy-tailed nature of model residuals was better captured with a Student’s t-distribution with a shared degrees-of-freedom parameter across proteins than with a Gaussian distribution. Consequently, using the Student’s t-distribution substantially improved the statistical calibration. Finally, we showed that variants predicted by AlphaMissense to be pathogenic, in contrast to the benign predictions, were enriched among underexpression outliers.

The full PROTRIDER model, which uses grid search to optimize the encoding dimension and explicitly incorporates relevant covariates and missingness modeling, consistently achieved the best performance, particularly when benchmarking for rare protein-disrupting variants on the mitochondrial disorder dataset. However, PROTRIDER (lite) remained a robust and computationally efficient alternative. Unlike the full version, PROTRIDER (lite) does not require hyperparameter tuning nor covariate specification, making it a convenient and perhaps more robust starting point. We therefore recommend running both versions. Enrichment for variants likely disrupting protein expression when available, or Q-Q-plots of tail probabilities and sample correlation heatmaps, can help decide whether the full model provided an improved fit to the data.

In this study, we presented the main results using the conservative Benjamini-Yekutieli procedure, which controls the FDR under arbitrary dependencies of the statistics. However, in our null model simulations, the FDR was also controlled using the Benjamini-Hochberg procedure, which assumes independence. Also, the enrichment for variants likely disrupting protein expression on the real data remained high, even though slightly reduced, with the Benjamini-Hochberg procedure. This suggests that the independence assumptions underlying the Benjamini-Hochberg method may not be substantially violated in practice, supporting its suitability in this context. Independent of the FDR procedure, using less stringent FDR cutoffs may be appropriate in scenarios where greater sensitivity to outliers or higher diagnostic rates is a priority. A further plausible scenario in rare disease diagnostics arises when genome analysis yields a candidate variant. Here, the nominal tail probability of the associated protein may be evaluated directly, without the need for FDR adjustment.

Given the enrichments we observed for rare and likely deleterious variants of various categories, we expect PROTRIDER to be of value for genetic diagnostic pipelines in addition to RNA-based analyses, especially to capture variant effects acting on the level of translation or protein stability. As we have shown earlier (Kopajtich et al., 2021) in the diagnostic context, proteomics can provide functional evidence for rare variants of uncertain significance. This is true for stop and frameshift variants but also for missense variants, which remain difficult variants to interpret. Here, we further showed a strong enrichment of AlphaMissense pathogenic variants in outliers detected at the protein level compared to other missense variants, highlighting a class of variants whose effects are often not captured by RNA-seq analyses.

The benchmarks presented in this study focus mostly on underexpression outliers. This is partly due to the nature of the available genomic information, as there were no copy number variations (CNVs) available for the mitochondrial datasets and only for some samples of the CRC65 panel, and partly because it is not straightforward to define classes of variants that lead to an increase in protein abundances. To benchmark overexpression outlier calls with genetic variants, further datasets with available copy number variants and larger sample sizes would be beneficial. Nevertheless, we could show using RNA-seq data that PROTRIDER (lite) overexpression calls were enriched for RNA-seq overexpression calls, indicating that the method can also capture overexpression outliers.

Missing values are common in mass spectrometry-based proteomics, especially with data-dependent acquisition (Ahlmann-Eltze and Anders 2019, Brenes et al. 2019, Karpievitch, Dabney and Smith 2012, Kong et al. 2022, Ahlmann-Eltze and Anders 2019), and often reflect low-abundance proteins (Kong et al. 2022). In PROTRIDER, incorporating the missingness binary mask allowed us to handle missing data more explicitly, rather than simply ignoring it in the loss function. This approach enabled the model to identify biologically relevant absences, such as protein intensities missing in certain samples due to regulation, sample-level, or batch-level technical dropouts. While PROTRIDER models missingness, we did not consider calling individual missing values as outliers. This could be relevant for non-TMT data-dependent acquisition proteomics but would require a substantial extension of the present study. In this context, future work may also investigate other methods such as variational autoencoders (Collier, Nazabal and Williams 2020, Nazábal et al. 2020) within the scope of outlier detection, particularly for incorporating probabilistic modeling and uncertainty estimates. However, the application of variational autoencoders to sample–protein level outlier detection would require substantial adaptation and evaluation of modeling choices, such as the design of an appropriate conditional prior and regularization terms to preserve reconstruction accuracy.

Furthermore, PROTRIDER only calls outliers at the protein level. However, entire protein complexes are often destabilized (Kremer et al. 2017, Kopajtich et al. 2021), therefore, additionally assessing outliers at the protein complex level could increase sensitivity. Additionally, peptide-level outliers may offer finer resolution, capturing variant-specific changes such as those caused by post-translational modifications or alternative splicing, which may have important functional consequences. At a more global level, one could also be interested in calling sample-level outliers, i.e., individuals whose entire proteome appears disrupted. In this case, multivariate outlier methods, including LSCP (Locally Selective Combination of Parallel Outlier Ensembles (Berger-Wolf and Chawla 2019)) and AnoGAN (Anomaly Detection with Generative Adversarial Networks (Schlegl et al. 2017)), could be worth being investigated. Another interesting extension of our work would be to model longitudinal datasets, which would allow for the detection of outlier values for individual subjects over time.

Although PROTRIDER was developed for proteomics data, it could also be applicable to other mass spectrometry-based measurements, such as metabolomics and lipidomics, which may share similar statistical properties. If the autoencoder-based correction and statistical calibration generalize well, PROTRIDER could support broader applications beyond proteomics.

## Data availability

No original data were created for this study. The RNA-seq and proteomics data of the mitochondrial dataset are available at GHGA (Dataset ID: <u>GHGAMCS41579681109068</u>).

The proteomics data of the NCI60 and CRC65 cell line panels were obtained through PRIDE accessions PXD013615 and PXD005354, as noted in (Frejno et al. 2020).

## Acknowledgments

We would like to thank the different users who have used preliminary PROTRIDER versions, as well as the members of the Gagneur lab who have given valuable feedback. We further thank the reviewers for their constructive and insightful comments during the revision process, which significantly helped us improve both the methods and the manuscript.

## Author contribution

Conceptualization: JG. Data Curation Management: IS, DKA, VAY. Formal Analysis: IS, DKA, GT, VAY. Resources: DS, HP. Software: IS, DKA, GT, SL. Supervision: VAY, JG. Validation: IS, DKA, VAY. Visualization: IS, DKA, GT, VAY, JG. Writing—original draft preparation: IS, DKA, VAY, JG. Writing—review & editing: all authors.

## Supplemental Figures

**Fig. S1:**
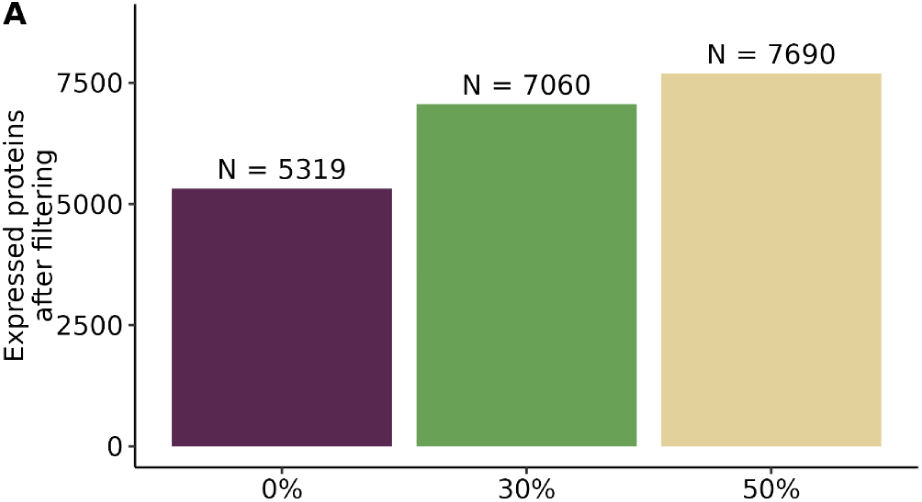
Expressed proteins and rare variants per category on the mitochondrial disorder dataset. **A**) Number of expressed proteins after filtering out proteins with too many missing values per protein, for three cutoffs on the maximal allowed percentage of missing values: no missing values (0%, purple), at most 30% missing values (green), and at most 50% missing values per protein (beige).

**Fig. S2:**
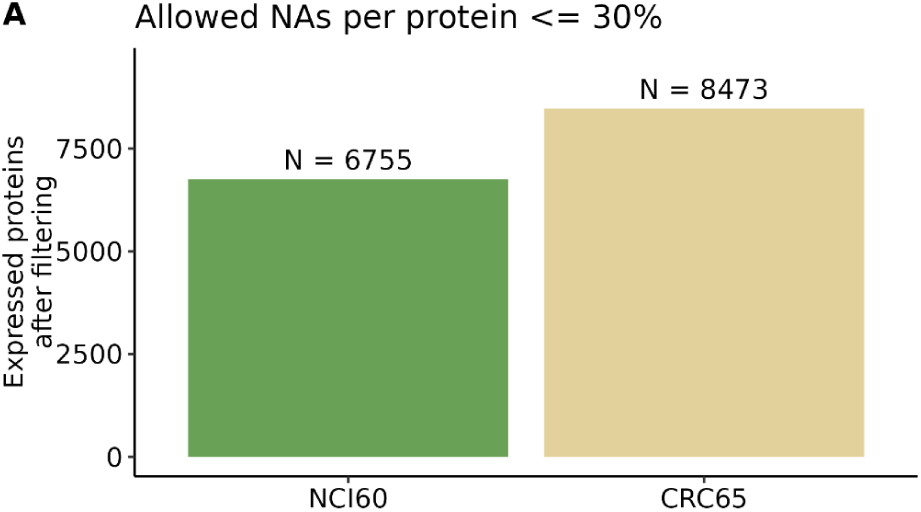
Expressed proteins and rare variants per category on the tumor cell line datasets. **A)** Number of expressed proteins after filtering out proteins with more than 30% missing values per protein, for the cell line panels NCI60 (green) and CRC65 (beige).

**Fig. S3:**
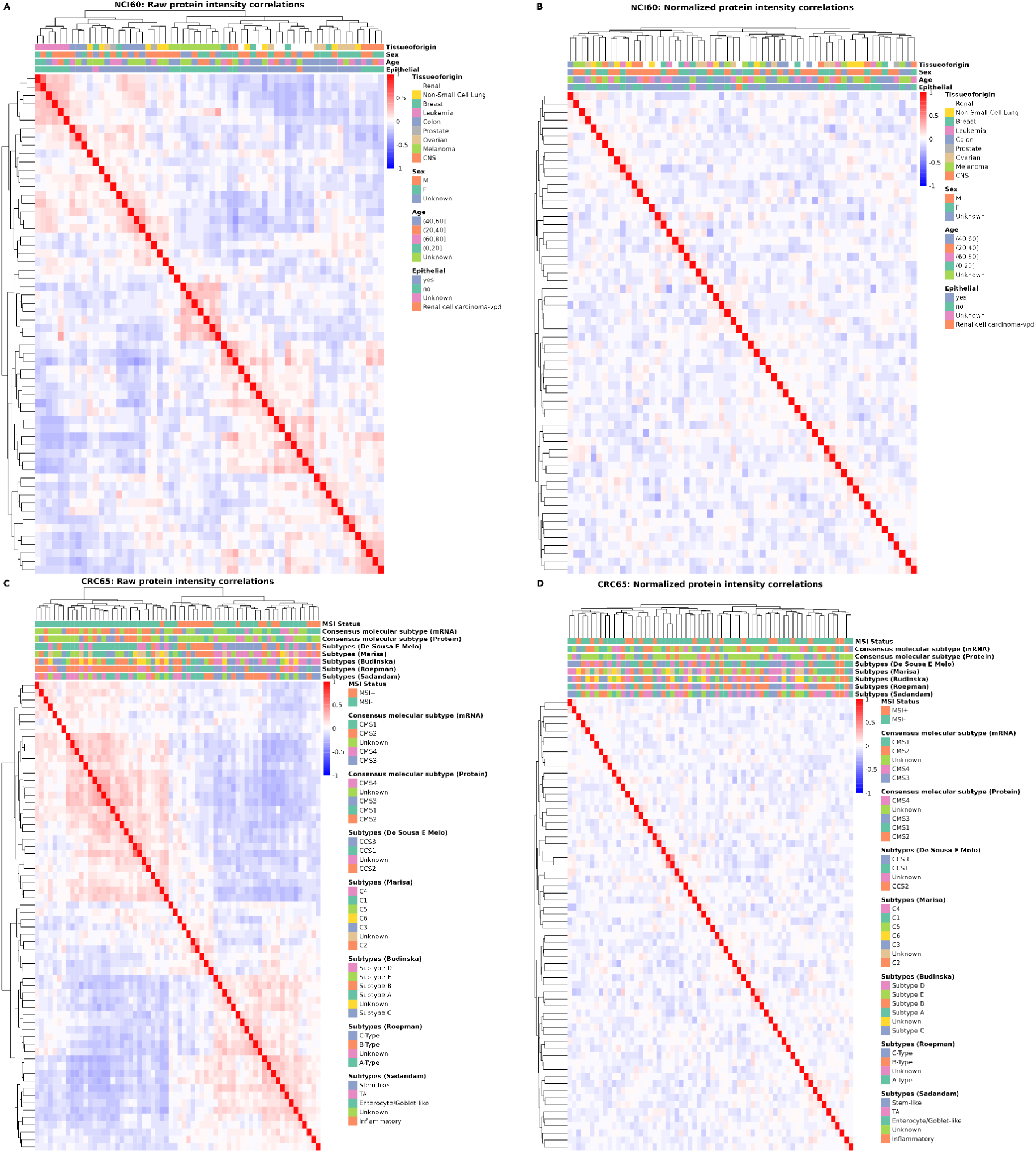
**Sample-sample correlations before and after autoencoder on the tumor cell line panels. A-B**) Heatmaps of sample-sample correlations of protein log-transformed intensities before (A) and after the PROTRIDER autoencoder correction for the PROTRIDER version using OHT for finding the optimal encoding dimension (B) on the NCI60 cell line panel data. **C-D)** Same as (A) and (B), respectively, but for samples from the cell line panel CRC65. The within-batch pairwise sample correlation was reduced from 0.15 ± 0.11 (mean ± standard deviation) to 0.079 ± 0.062 for NCI60 and from 0.11 ± 0.074 to 0.05 ± 0.04 for CRC65’

**Fig. S4:**
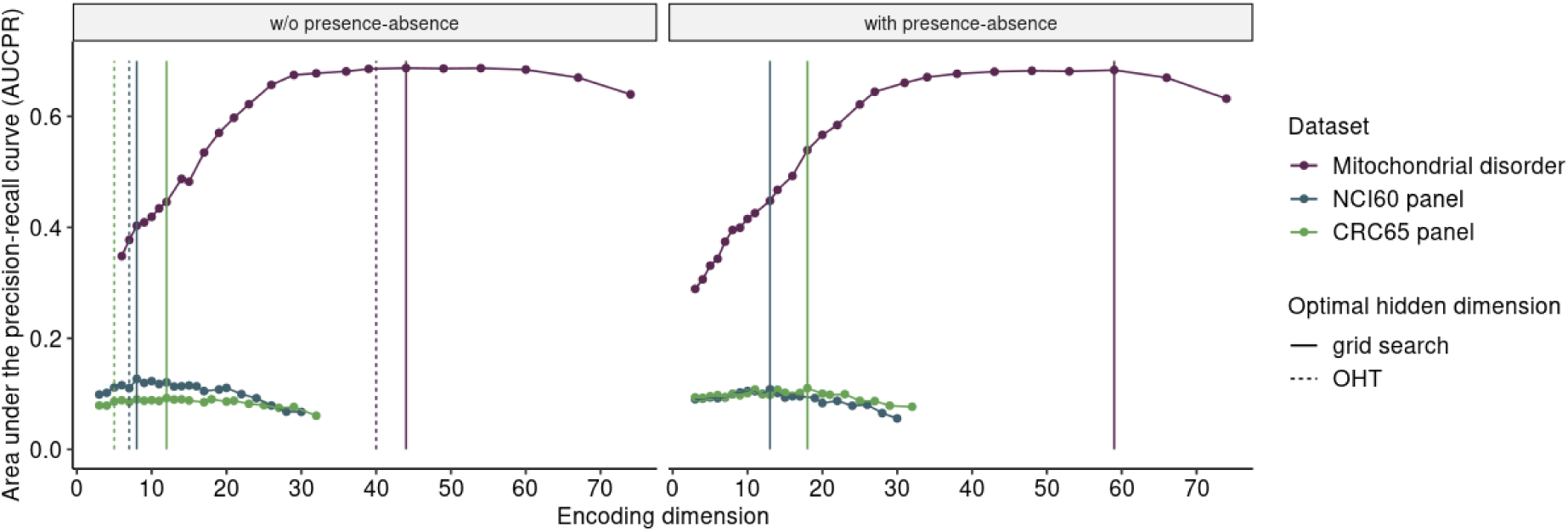
Optimal encoding dimensions found by PROTRIDER and PROTRIDER (lite). The obtained area under the precision-recall curve of recovering injected outliers is plotted against the candidate encoding dimensions for the PROTRIDER approach without (left) and with (right) missingness modelling, i.e. presence-absence information. The vertical lines indicate the optimal encoding dimension found for each dataset with the grid search approach and the OHT approach.

**Fig. S5:**
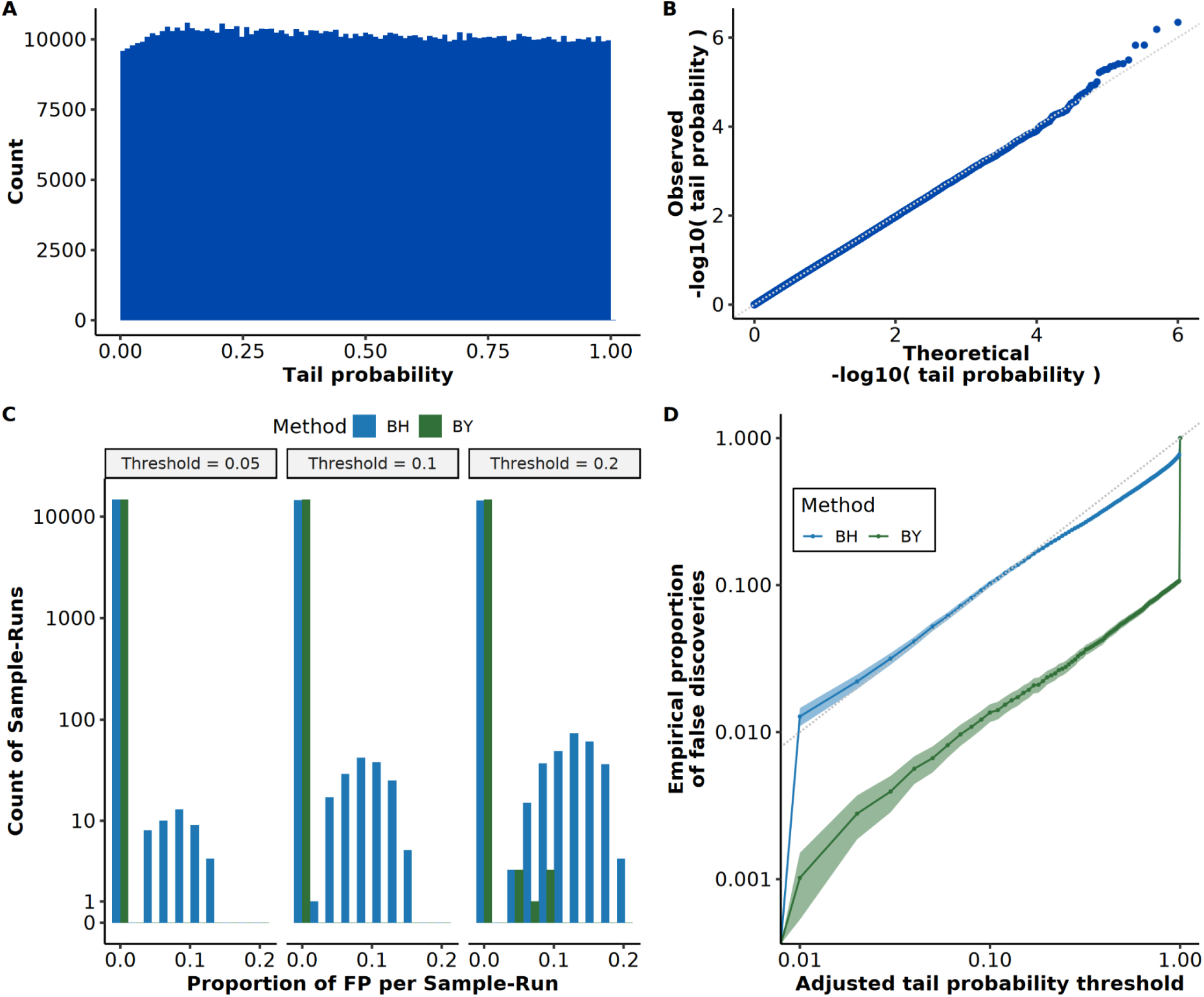
Empirical FDR validation with PROTRIDER on simulated data under the null model. **A)** Histogram showing the distribution of two-sided tail probabilities computed by PROTRIDER fitted on a simulated intensity matrix under the assumption that model residuals follow a Student’s t-distribution. **B)** Quantile-Quantile plot comparing the observed-log10(tail probabilities) to their theoretical quantiles under the null (uniform distribution). **C)** Histograms showing the proportions of false positives across simulated samples, stratified by significance threshold on adjusted tail probabilities with the method of Benhamini-Hochberg (BH, blue) and Benjamini-Yekutieli (BY, green). **D)** Adjusted tail probability thresholds (x-axis) against empirically observed proportions of false discoveries (y-axis) with the method of Benhamini-Hochberg (BH, blue) and Benjamini–Yekutieli (BY, green), estimated from 100 null simulations using residuals sampled from a fitted Student’s t-distribution. The diagonal dashed line indicates perfect FDR calibration. Ribbons mark 95% confidence intervals.

**Fig. S6:**
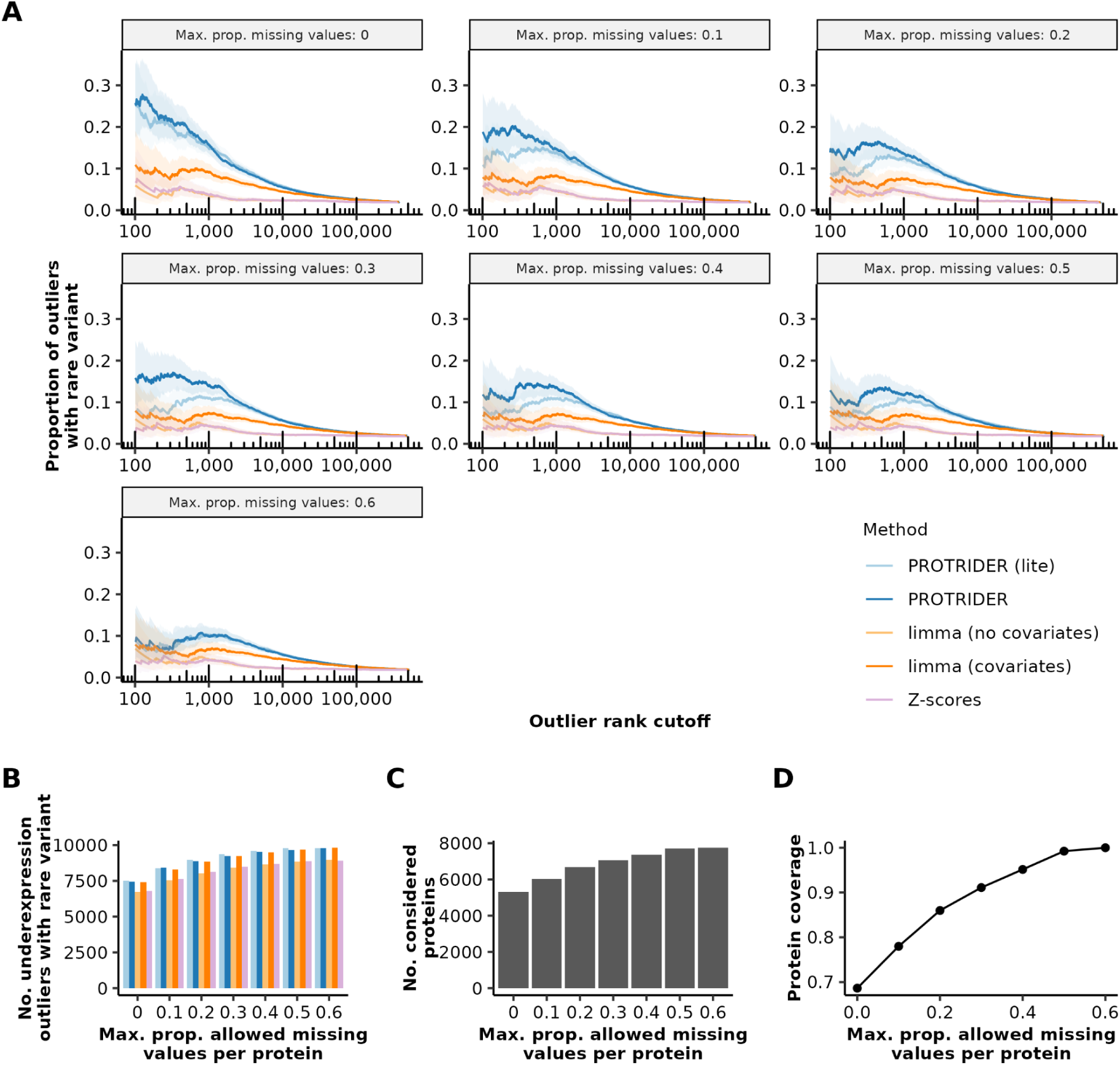
PROTRIDER performance on the mitochondrial disorder dataset for different missing value thresholds. A, Proportion of outliers with at least one rare variant likely disrupting protein expression on the mitochondrial disorder dataset for underexpression outliers calls from PROTRIDER (blue), PROTRIDER (lite, light blue), the limma-based method with covariates (orange) and without covariates (light orange), and the Z-score-based method (violet) for different thresholds for removing proteins with more than the specified the maximal proportion of missing values per protein. Ribbons mark 95% confidence intervals. **B,** Total number of underexpression outliers with a rare variant likely disrupting protein expression for the different missing value thresholds. **C,** Total number of considered proteins for the different missing value thresholds. **D,** Protein coverage, i.e,. number of proteins considered over the total number of proteins in the mitochondrial disorder dataset, for the different missing value thresholds.

**Fig. S7.**
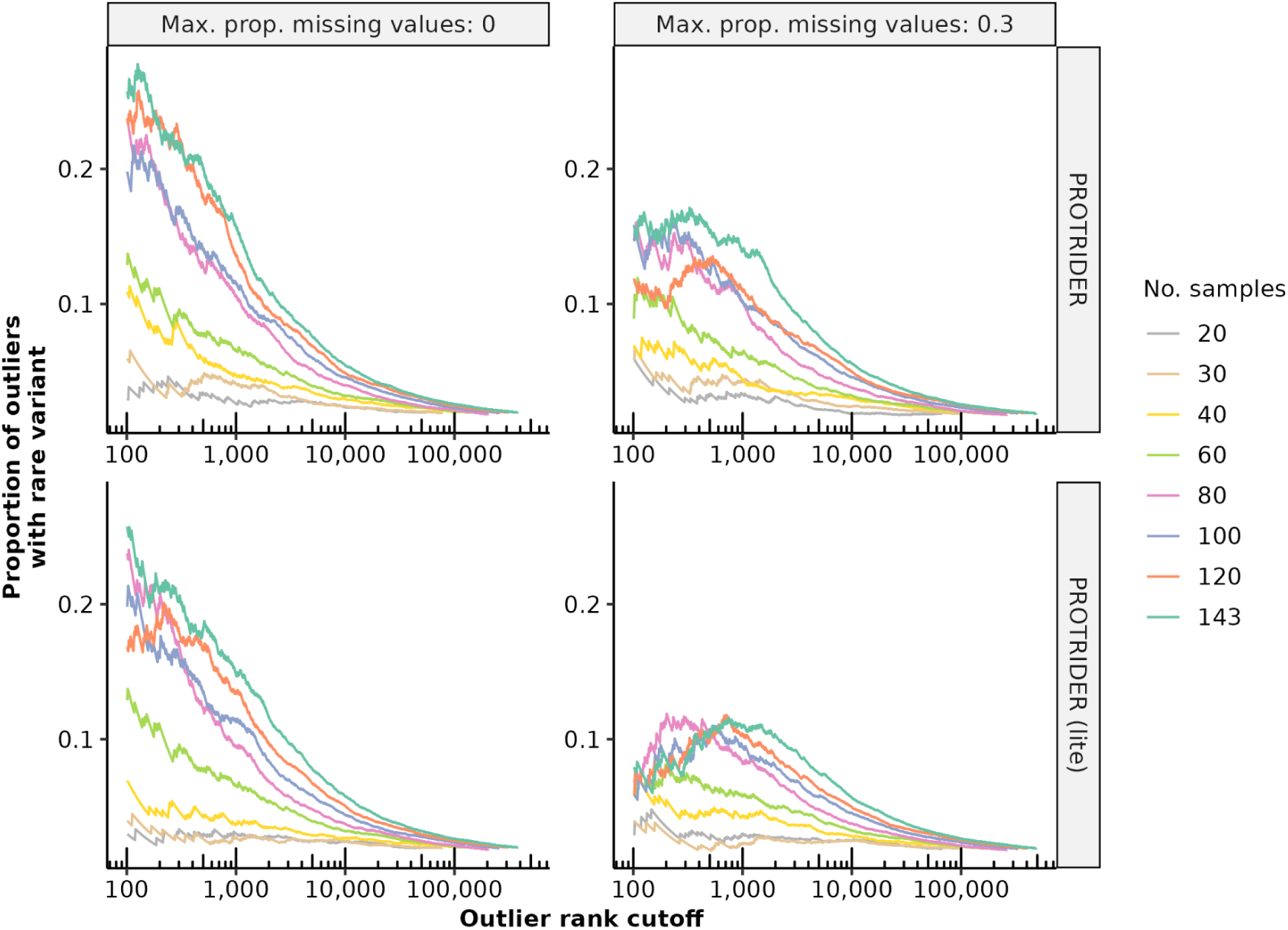
PROTRIDER performance on random subsets of the mitochondrial disorder dataset. Proportion of outliers with at least one rare variant likely disrupting protein expression on the mitochondrial disorder dataset for underexpression outliers calls from PROTRIDER and PROTRIDER (lite) for different sample sizes obtained after randomly subsetting the mitochondrial disorder dataset and for two different missing value thresholds (0 and 0.3) for the maximal proportion of missing values per protein.

**Fig. S8.**
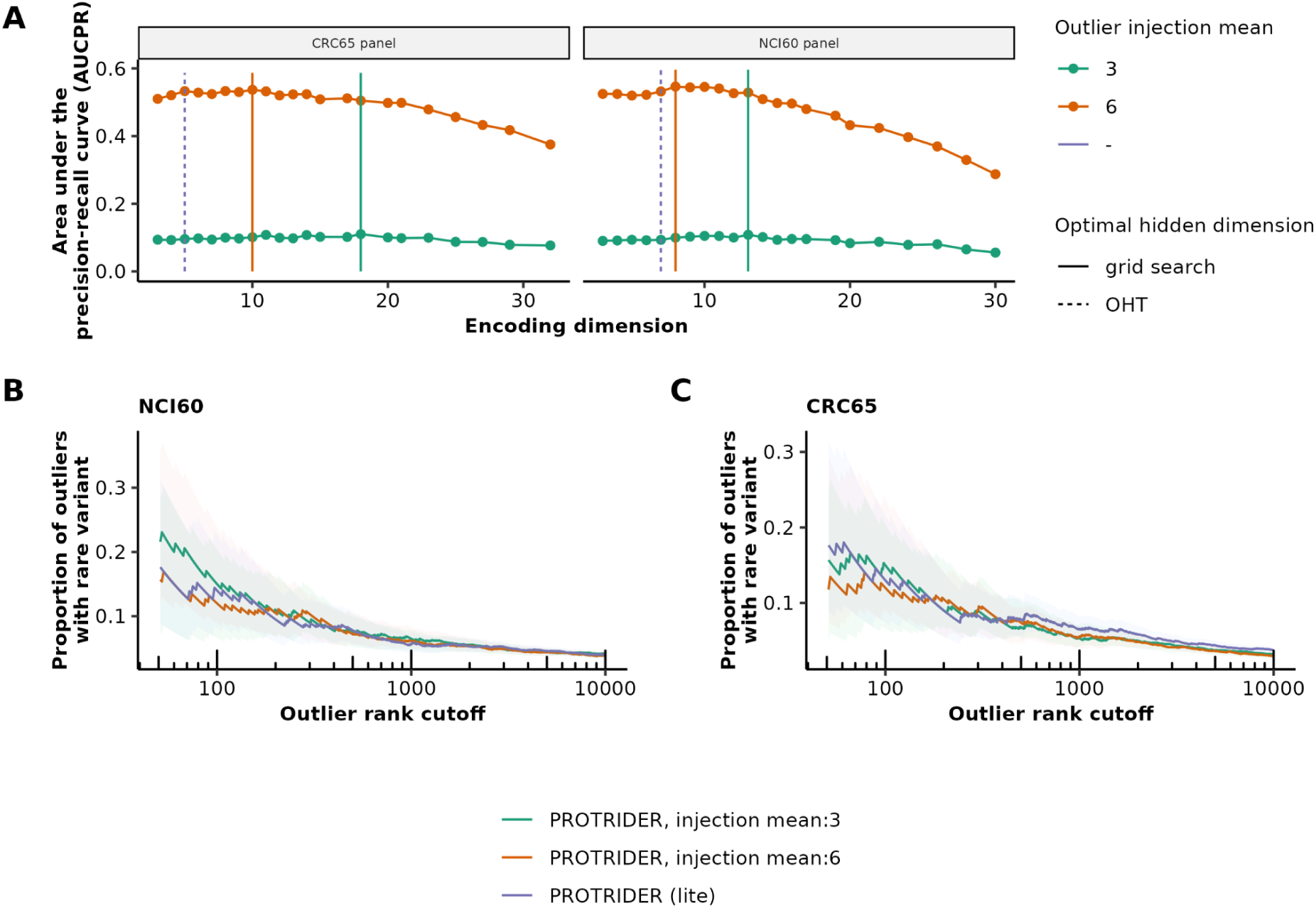
PROTRIDER performance on the tumor cell line panels for different outlier injection settings. A, The obtained area under the precision-recall curve of recovering injected outliers with injection mean 3 (green) and 6 (orange) is plotted against the candidate encoding dimensions for PROTRIDER. The vertical lines indicate the optimal encoding dimension found for each dataset with the grid search approach and the OHT approach. **B,** Proportion of outliers with at least one rare variant likely disrupting protein expression on the mitochondrial disorder dataset for underexpression outlier calls from PROTRIDER based on an injection mean of 3 (green), 6 (orange), and PROTRIDER (lite, violet) for the tumor cell line panel NCI60. **C**, Same as for,B but for the cell line panel CRC65

**Fig. S9:**
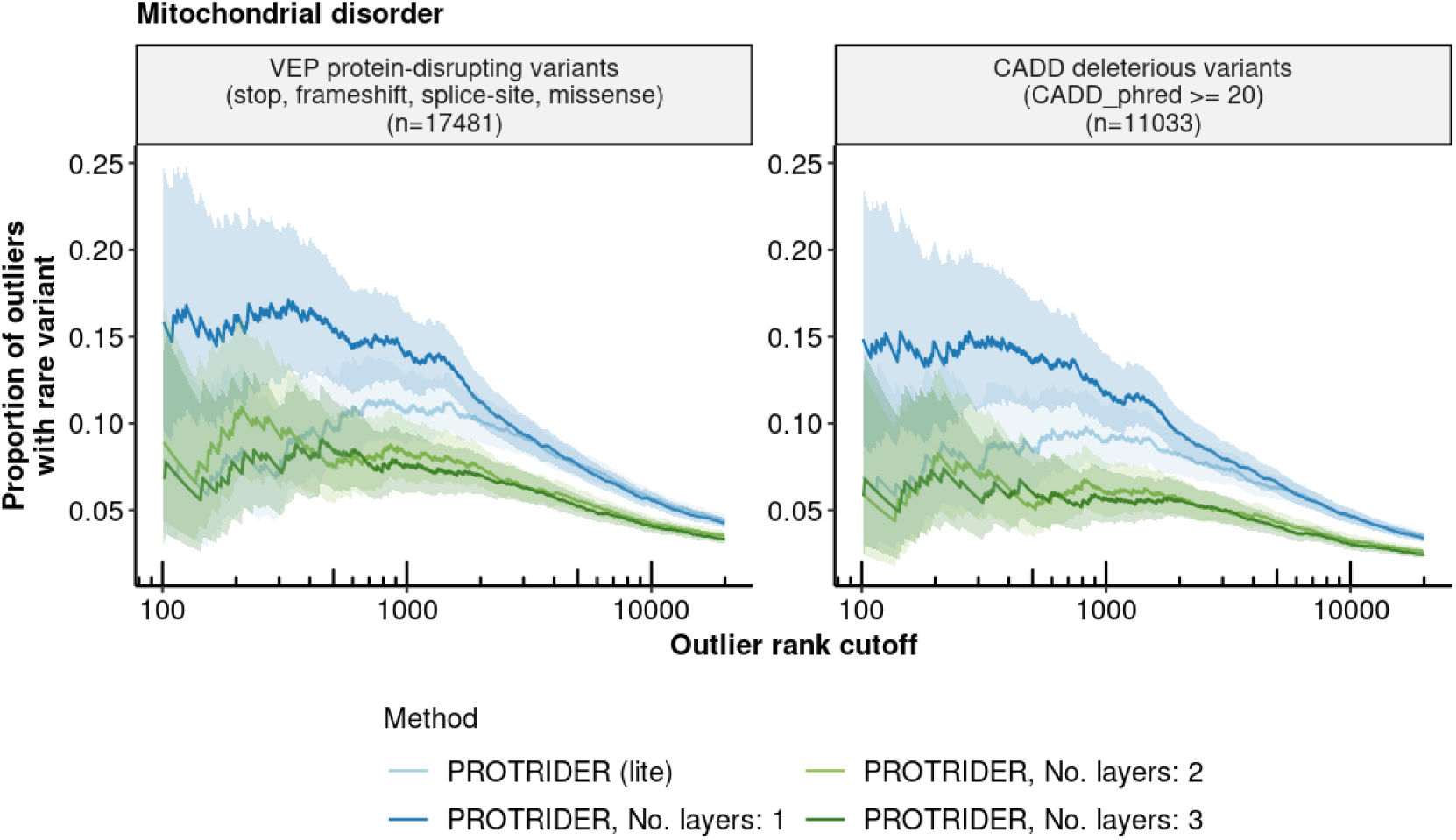
Linear autoencoder model outperforms multi-layer autoencoder models. Proportion of outliers with at least one rare variant on the mitochondrial disorder dataset for underexpression outliers calls from PROTRIDER with different number of layers and, PROTRIDER (lite) with two sets of rare variant categories as ground truth proxies: i) VEP stop, frameshift, direct split-site, and missense variants and ii) CADD deleterious variants (PHRED score ≥ 20).

**Fig. S10:**
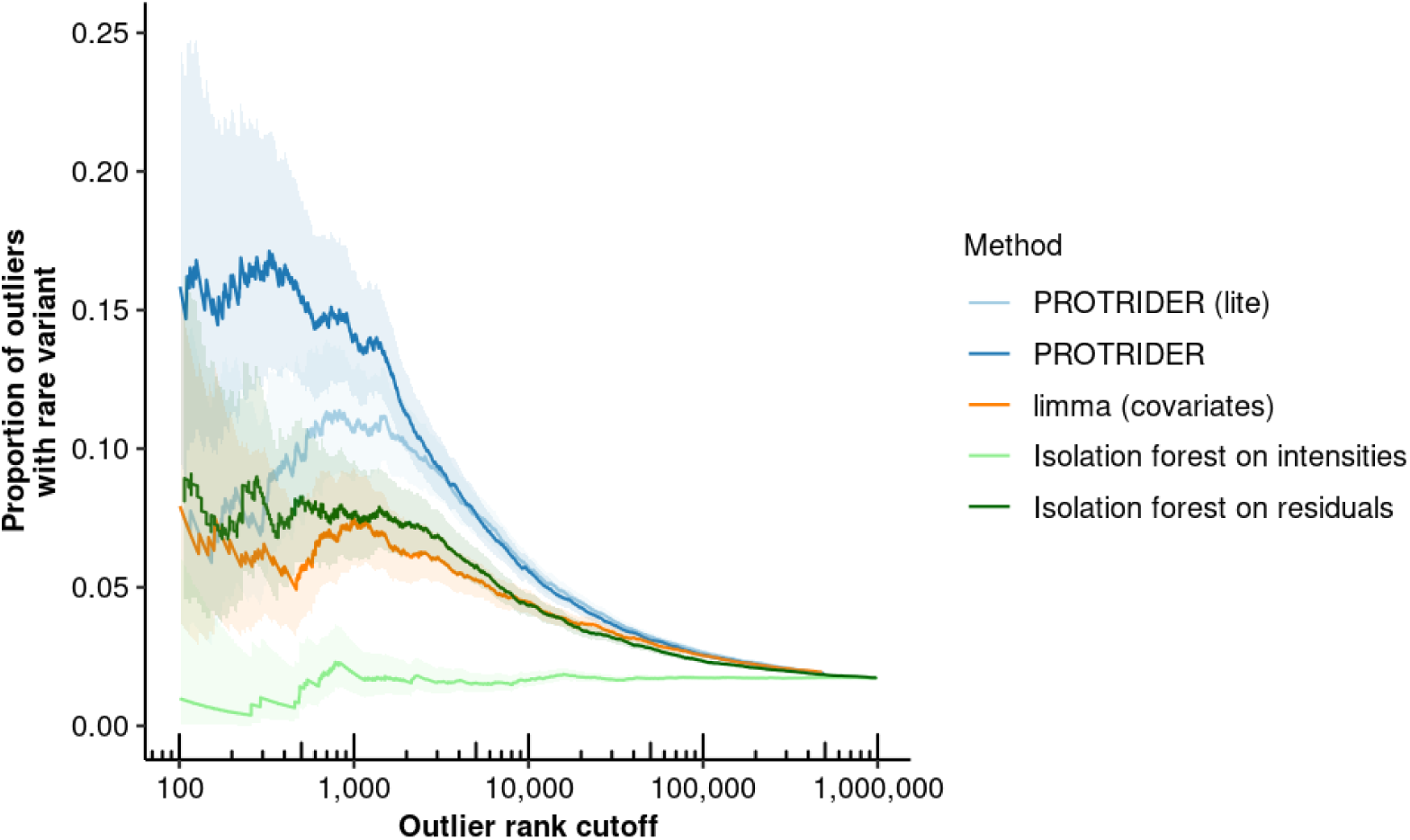
PROTRIDER performance compared to an alternative approach based on Isolation forests. Proportion of outliers with at least one rare variant likely disrupting protein expression on the mitochondrial disorder dataset for underexpression outliers calls from PROTRIDER (blue), PROTRIDER (lite, light blue), the limma-based method with covariates (orange), a protein-specific Isolation forest fitted on preprocessed intensities (light green), and fitted on the residuals returned by PROTRIDER (dark green). Ribbons mark 95% confidence intervals.

**Fig. S11:**
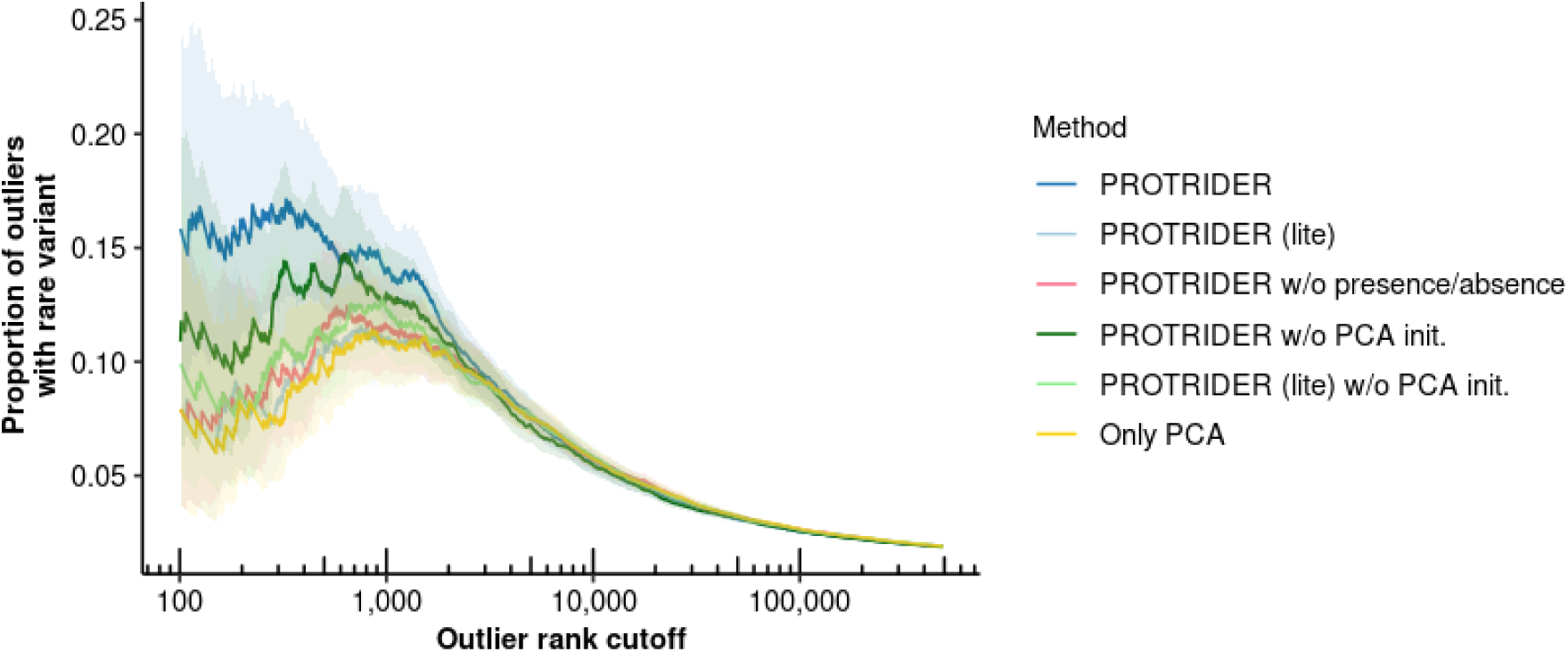
PROTRIDER ablation study. Proportion of outliers with at least one rare VEP stop, frameshift, direct split-site, and missense variant on the mitochondrial disorder dataset for underexpression outlier calls from PROTRIDER (dark blue), PROTRIDER (lite, light blue), PROTRIDER without missingness modeling (rose), PROTRIDER without PCA initialization (light green and dark green), and only PCA projection with the number of principal components derived from the OHT procedure.

**Fig. S12:**
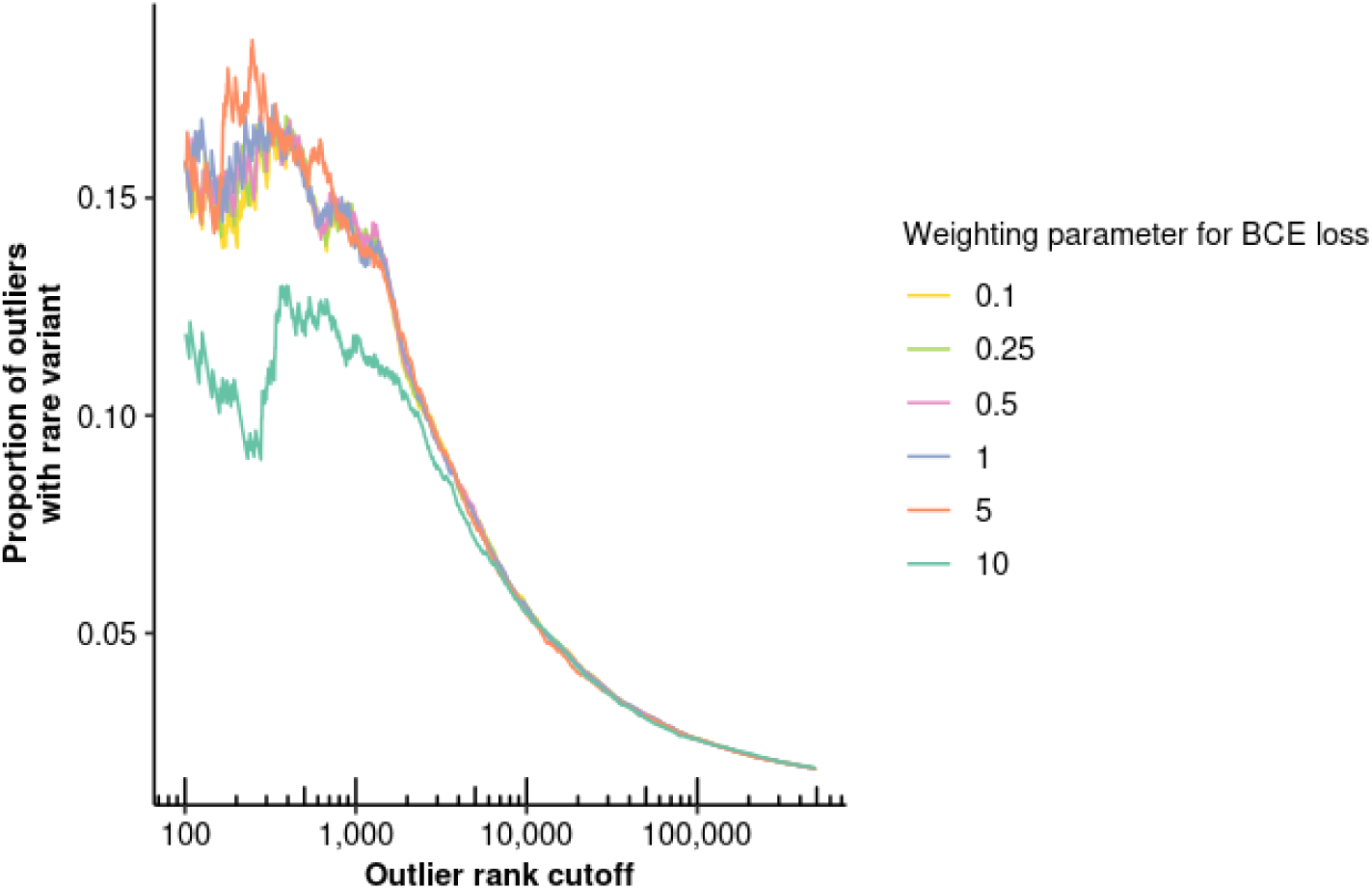
Impact of weighting factor for binary cross entropy compared to regression loss. Proportion of outliers with at least one rare VEP stop, frameshift, direct split-site, and missense variant on the mitochondrial disorder dataset for underexpression outlier calls from PROTRIDER fitted with different weighting factors to aggregate the binary cross entropy and the mean squared error loss to optimize the prediction of protein intensities and presence probabilities.

**Fig. S13:**
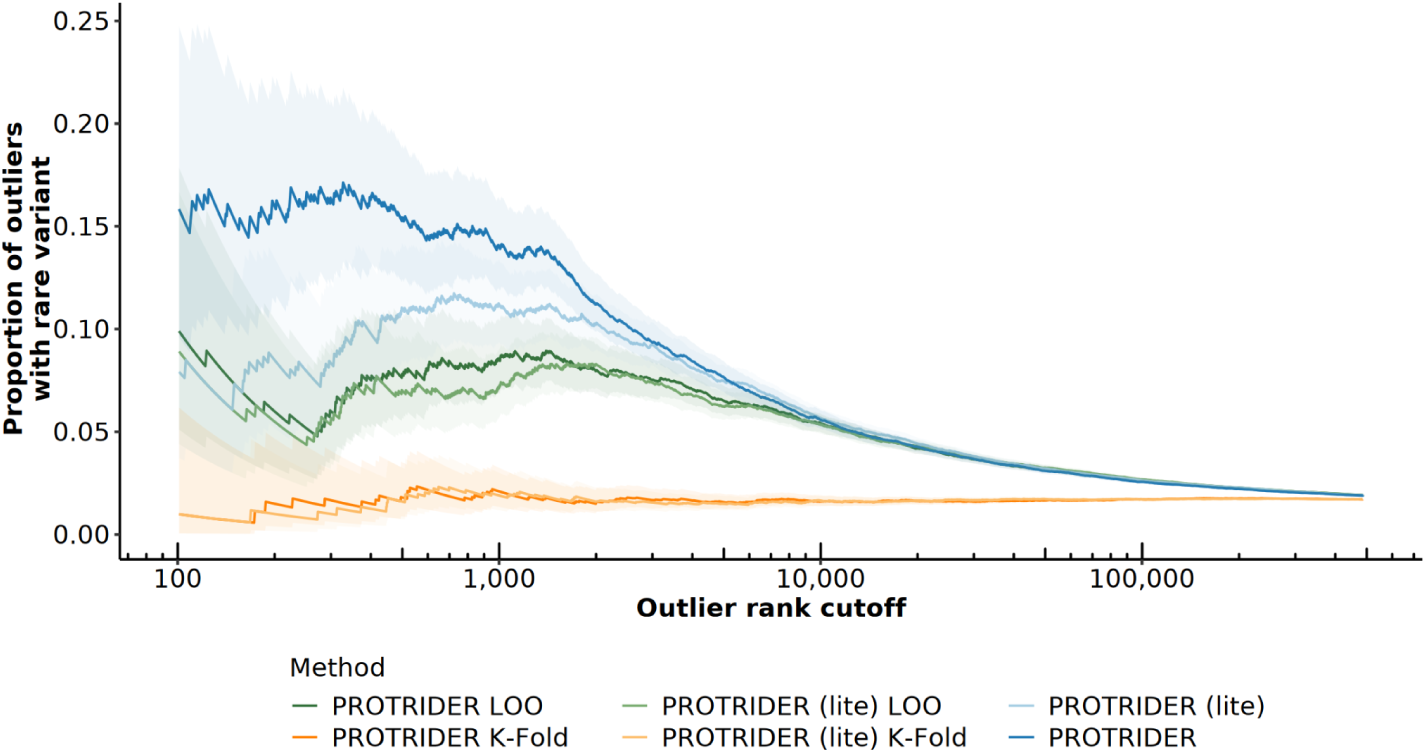
Models trained on the full dataset perform best compared to cross-validation approaches. Proportion of underexpression outliers with at least one rare variant in the mitochondrial disorder dataset. Variants include VEP stop, frameshift, direct splice-site, and missense variants, used as ground truth proxies. The following models are compared: PROTRIDER (lite) with leave-one-out cross-validation (light green), *k*-fold cross-validation (light orange), and no cross-validation (light blue); PROTRIDER with leave-one-out cross-validation (green), *k*-fold cross-validation (orange), and no cross-validation (blue). Ribbons mark 95% confidence intervals.

**Fig. S14:**
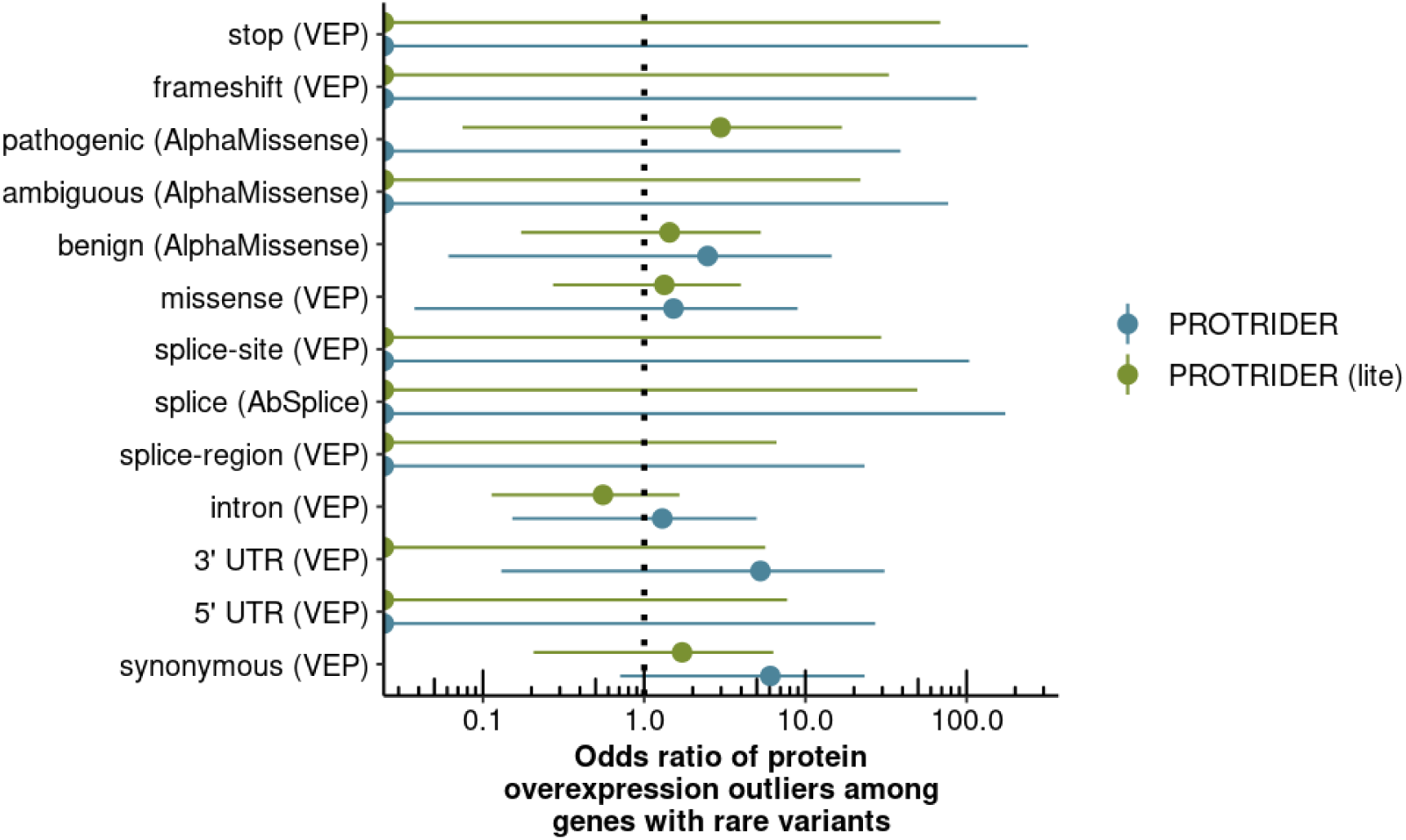
Genetic determinants of protein overexpression outliers. Odds ratios and their 95% confidence intervals (Fisher’s test) of the enrichment of the proportion for each variant category among overexpression outliers compared to the background proportion of the non-outliers.

**Fig. S15:**
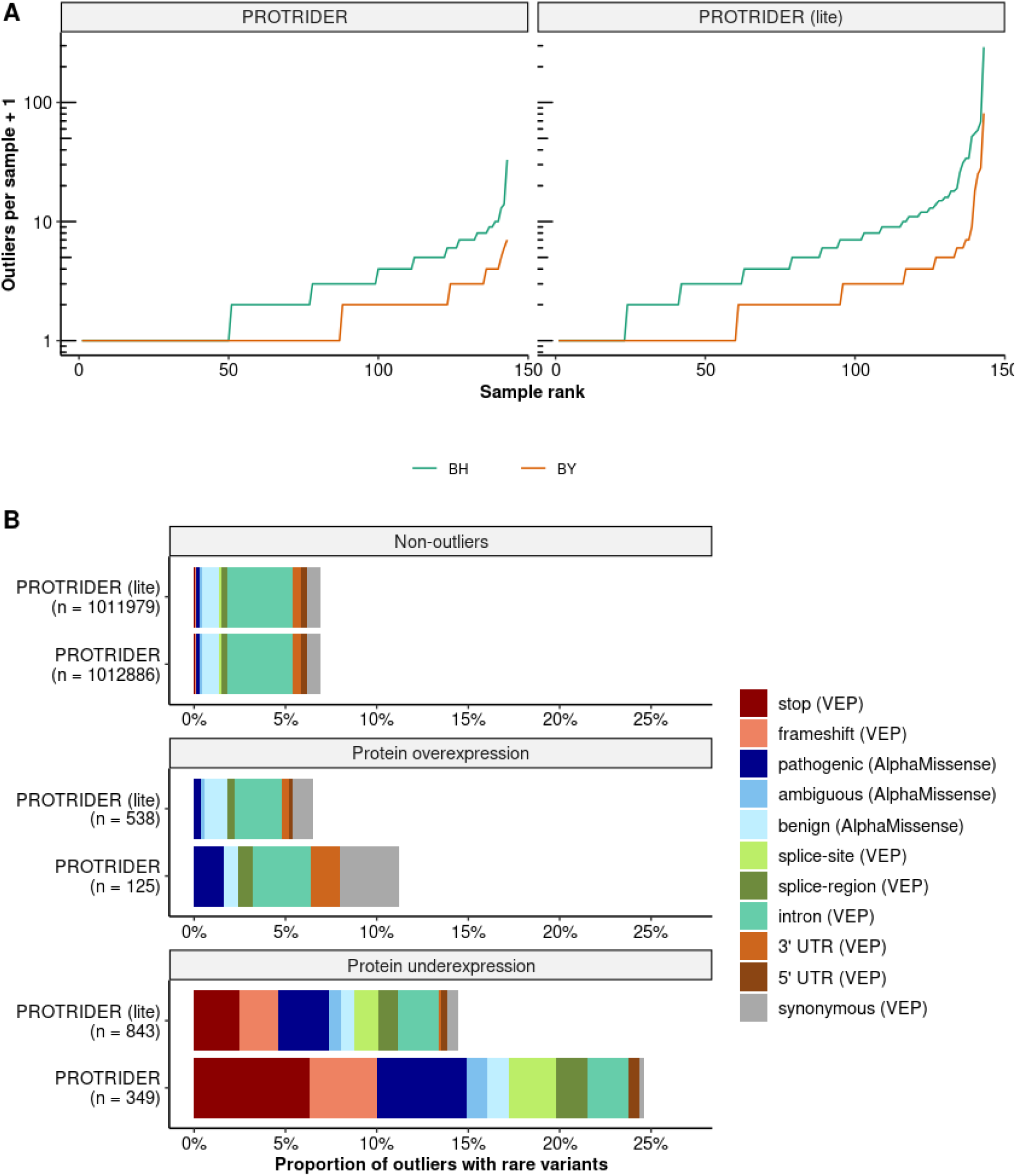
Method comparison for tail probability adjustment. A, Sorted number of protein abundance outliers per sample obtained after adjusting the obtained tail probabilities with the methods of Benjamini and Yekutieli (BY) and Benjamini and Hochberg (BH) for the two PROTRIDER versions (facets). **B**, Same as Fig. 4A, but with the BH procedure for controlling the false discovery rate.

